# Enhanced antifungal activity of NK-92 cells against *Candida albicans* mediated by a mannan-specific chimeric antigen receptor

**DOI:** 10.1101/2024.11.18.623666

**Authors:** Gabriela Yamazaki de Campos, Júlia Garcia Guimarães, Michele Procópio Machado, Patrícia Kellen Martins Oliveira Brito, Ben Shin, Antonio Di Maio, Douglas dos Santos, Patricia Vianna Bonini Palma, Thaila Fernanda dos Reis, Gustavo Henrique Goldman, Angelina S. Palma, Steve J. Matthews, Ten Feizi, Yan Liu, Thiago Aparecido da Silva

## Abstract

Chimeric antigen receptors (CARs) offer promising prospects for innovative cell-based therapies against invasive fungal infections such as invasive candidiasis. Here, we have developed four CARs targeting *Candida albicans* with distinct single-chain variable fragments (scFvs): scFv3-CAR, scFv5-CAR, scFv12-CAR, and scFvκ3-1-CAR. In T cells, scFv5-CAR induced IL-2 expression in response to *C. albicans* hyphae, while scFv3-CAR and scFv12-CAR did not mediate cell activation against *C. albicans*. Notably, scFvκ3-1-CAR mediated the strongest cell activation against *C. albicans* yeast, hyphae, and other clinically relevant *Candida* species. scFvκ3-1-CAR-NK-92 cells exhibited elevated IFN-γ and CD107a expression, reducing *C. albicans* viability. NOD scid gamma (NSG) mice treated with scFvκ3-1-CAR-NK-92 cells had reduced *C. albicans* burden in the kidney 24 hours postinfection. We showed that scFvκ3-1-CAR targets *C. albicans* mannan but no other glycans in glycan microarray screening analyses. These findings reveal the scFvκ3-1-CAR potential as a therapeutic strategy for treating *Candida* spp. by modifying peripheral blood mononuclear cells.

**Importance:** Recent studies on novel immunotherapies, including chimeric antigen receptor (CAR)-T cells, have shown promising results in preclinical models against invasive fungal infections (IFIs). However, the application of CAR technology in natural killer (NK) cells for treating IFIs remains unexplored. NK cells play a key role in early fungal clearance due to their antifungal activity mediated by granzymes, perforins, and the secretion of proinflammatory cytokines. This study is the first to demonstrate the feasibility and efficacy of CAR-modified NK cells targeting *Candida* spp. We provided proof-of-concept data showing that CAR-expressing NK cells exhibit enhanced activation and antifungal effects against clinically relevant *Candida* species by targeting mannan in the fungal cell wall. These findings are significant as they open new avenues for developing CAR-NK-based therapies to treat invasive candidiasis – a severe infection with limited treatment options and high mortality rates, particularly in immunocompromised patients.

## Introduction

*Candida* spp. are commensal organisms that reside in the gastrointestinal tract and on the skin and can cause invasive candidiasis by spreading from their natural colonization sites to the bloodstream, leading to systemic dissemination (1). Approximately 1,565,000 individuals are affected by candidemia or invasive candidiasis annually, resulting in 995,000 deaths (63.6%) (2). The use of antibiotics combined with central venous catheterization, total parenteral nutrition, dialysis, and extensive abdominal surgery are major primary risk factors for invasive candidiasis (1). Although *Candida albicans* remains the predominant species responsible for these infections, other species, such as *Candida glabrata*, *Candida tropicalis*, *Candida parapsilosis,* and *Candida krusei,* also contribute to the occurrence of candidiasis (1, 3, 4). *Candida auris* has emerged globally as a multidrug-resistant fungus and has been associated with outbreaks in intensive care units, including among COVID-19 patients, acting as a secondary infection with a mortality rate of approximately 50% (5, 6).

Various microenvironmental factors in the host induce the transition from the yeast to the hyphal form of *C. albicans*, which involves the regulation of genes encoding proteins and glycosyltransferases involved in the cell wall composition, facilitating the evasion of *Candida* spp. from the host immune response (7–9). The cell wall of *C. albicans* is composed of 80-90% carbohydrates. The outer layer is composed mainly of N- and O-linked mannose polymers, while the inner layer is predominantly composed of β-1,3-glucan and β-1,6-glucan (10). The carbohydrate components of the *C. albicans* cell wall are pathogen-associated molecular patterns (PAMPs) that are recognized by innate immune cells (11, 12). The C-type lectin receptors, Toll-like receptors (TLRs) 2 and 4, and the mannose receptor recognize polysaccharides on the surface of *C. albicans* yeast and hyphae, initiating the activation of the innate immune cells, including natural killer (NK) cells (13–15). NK cells play a crucial role in controlling *C. albicans* filament growth in the initial hours of infection, which is achieved through the release of cytotoxic granules such as perforins and granzymes (16). Additionally, NK cells release IFN-γ, TNF-α, GM-CSF, and chemokines for NK cell migration, and IFN-γ production activates polymorphonuclear leukocytes, increasing antifungal activity and promoting T-cell differentiation (17). The differentiation of CD4^+^ T-helper (Th) and CD8^+^ T-cytotoxic (Tc) cells is crucial for inhibiting the growth of *C. albicans* hyphae *in vitro* (18), with the Th1, Th17, Tc1, and Tc17 cell subsets playing active roles in protecting against invasive candidiasis (19, 20). The cytotoxic activity of CD8^+^ T cells, which involves cytolytic granules and proinflammatory mediators, is necessary for combating invasive candidiasis (21). Failure of these cells to drive adaptative immune responses to *C. albicans* may be associated with structural remodeling of the *C. albicans* cell wall (22). Moreover, *C. albicans* can inhibit phagolysosome maturation and induce macrophages to transition from an M1 phenotype to a less inflammatory M2 macrophage subset (23, 24).

Although antifungal drugs are the primary clinical treatment for invasive candidiasis, the prevalence of microbial resistance and associated side effects highlight the necessity for alternative and complementary therapeutic modalities (25, 26). Immunotherapy approaches have been evaluated for treating invasive fungal infections (IFIs), and adoptive T- and NK-cell therapy shows potential for clinical translation (27). In this context, chimeric antigen receptor (CAR) technology for targeting IFIs has advanced and arisen as a potential strategy, as demonstrated by previous studies (28–33). CARs are receptors designed to be expressed by immune cells, redirecting them to interact with a specific antigen (protein, glycoprotein, carbohydrates, lipids, and others) localized on the cell surface. The antigen-binding domain of a CAR consists of a single-chain variable fragment (scFv) derived from a monoclonal antibody specific to the target antigen (34). CARs also include a transmembrane domain, often derived from CD28 or CD8 molecules, which anchors the receptor to the cell surface (35). The signaling domain of a second-generation CAR is composed of activation motifs from the CD3ζ chain coupled with a costimulatory molecule (such as CD28, 4-1BB, or iCOS)(34). Over the past decade, human T cells have been engineered with CARs to target *Aspergillus fumigatus* conidia and/or hyphae, resulting in the control of conidium-hyphal switching both *in vitro* and *in vivo* (32). Additionally, CAR-T cells targeting *A. fumigatus* hyphae conferred antifungal efficacy and improved the survival of mice (33). Our group developed a CAR to target glucuronoxylomannan (GXM) expressed in *Cryptococcus* spp. capsules, named GXMR-CAR (29, 30, 36). We found that GXMR-CAR-T cells recognized titan cells of *Cryptococcus gattii* and *Cryptococcus neoformans* (29) and reduced the percentage of *C. neoformans* titan cells in mouse lung tissue (30). To date, NK cells have not been explored for CAR-based therapy against IFIs. However, there has been notable interest in NK cells for cancer therapies, particularly solid tumors (37, 38). The NK-92 cell line has been used in clinical trials involving CAR-based treatments, offering the advantage of being an off-the-shelf cell therapy product (39). This highlights their potential in adoptive cell therapy and suggests that similar approaches could be adapted for the treatment of IFIs.

scFvs are typically utilized to design ligand-binding domains for novel CAR constructs. Several scFvs have been thoroughly investigated for their ability to bind to specific antigens of *C. albicans.* In this study, we selected four scFvs derived from human monoclonal antibody fragments generated by phage display and previously described in the literature (40–42) to design novel CARs. scFv3 (42), scFv5 (41), and scFv12 (41) were described to target *C. albicans* hyphae, and scFv κ3-1 (40) targets both yeast and hyphae. scFv3 recognizes the adhesin Als3p, a protein associated with germ tube and hyphal formation (42), while scFv5 likely recognizes the product of the gene *SPT6* (43). The specific antigens for scFv12 and scFv κ3-1 remain unidentified, although it was indicated that scFv12 binds to a protein (41), while scFv κ3-1 binds to a carbohydrate (40).

To further advance the development of CAR technology for targeting fungi, CARs capable of redirecting NK-92 cells to recognize *Candida* species were developed in the current study. Here, four second-generation CAR constructs containing distinct scFvs were designed (scFv3-, scFv5-, scFv12-, and scFvκ3-1-CAR). Jurkat and NK-92 cells were separately bioengineered with CARs, and scFvκ3-1-CAR mediated the strongest cell activation against *C. albicans* yeast and hyphae, as demonstrated by the high production of proinflammatory cytokines and activation markers. Additionally, scFvκ3-1-CAR-Jurkat cells were activated in the presence of *C. glabrata, C. tropicalis,* and *C. auris*. *In vitro* studies revealed that scFvκ3-1-CAR-NK-92 cells increased fungal damage by releasing cytotoxic granules, and the infusion of scFvκ3-1-CAR-NK-92 cells in NSG mice resulted in a significant reduction in the fungal burden in the kidneys. Finally, a glycan microarray demonstrated that scFvκ3-1-CAR specifically targets the mannan of *Candida* spp. Thus, this study pioneers the bioengineering of CAR-NK-92 cells to target and damage fungal species, providing proof-of-concept that the scFvκ3-1-CAR construct could be used in future studies to enhance the fungicidal activity of human primary T and NK cells against *Candida* species.

## Results

### *C. albicans*-specific CAR constructs generated from distinct scFvs specific to yeast and/or hyphal forms

*C. albicans*-specific CAR constructs were designed with different scFvs previously characterized to recognize the yeast and hyphae of *C. albicans* (40–42), and chimeric receptors, namely, scFv3-CAR, scFv5-CAR, scFv12-CAR, and scFvκ3-1-CAR, were generated. These CAR constructs are schematically represented in Figure 1a. The CD8 molecule is the hinge/transmembrane domain, followed by the intracellular domain comprising the activation motifs of CD137 and CD3ζ molecules. The enhanced green fluorescent protein (EGFP) serves as a marker. The coding sequences of the *C. albicans*-specific CAR constructs were cotransfected with pMD2.G and psPAX2 accessory plasmids into HEK-293T cells. The titers of the scFv3-CAR, scFv5-CAR, scFv12-CAR, and scFvκ3-1-CAR lentiviral particles were determined in the Jurkat cell line, which exhibited high titers of lentiviral particles (Fig. 1b), whereas the scFvκ3-1-CAR construct exhibited a lower titer.

**Figure 1.**
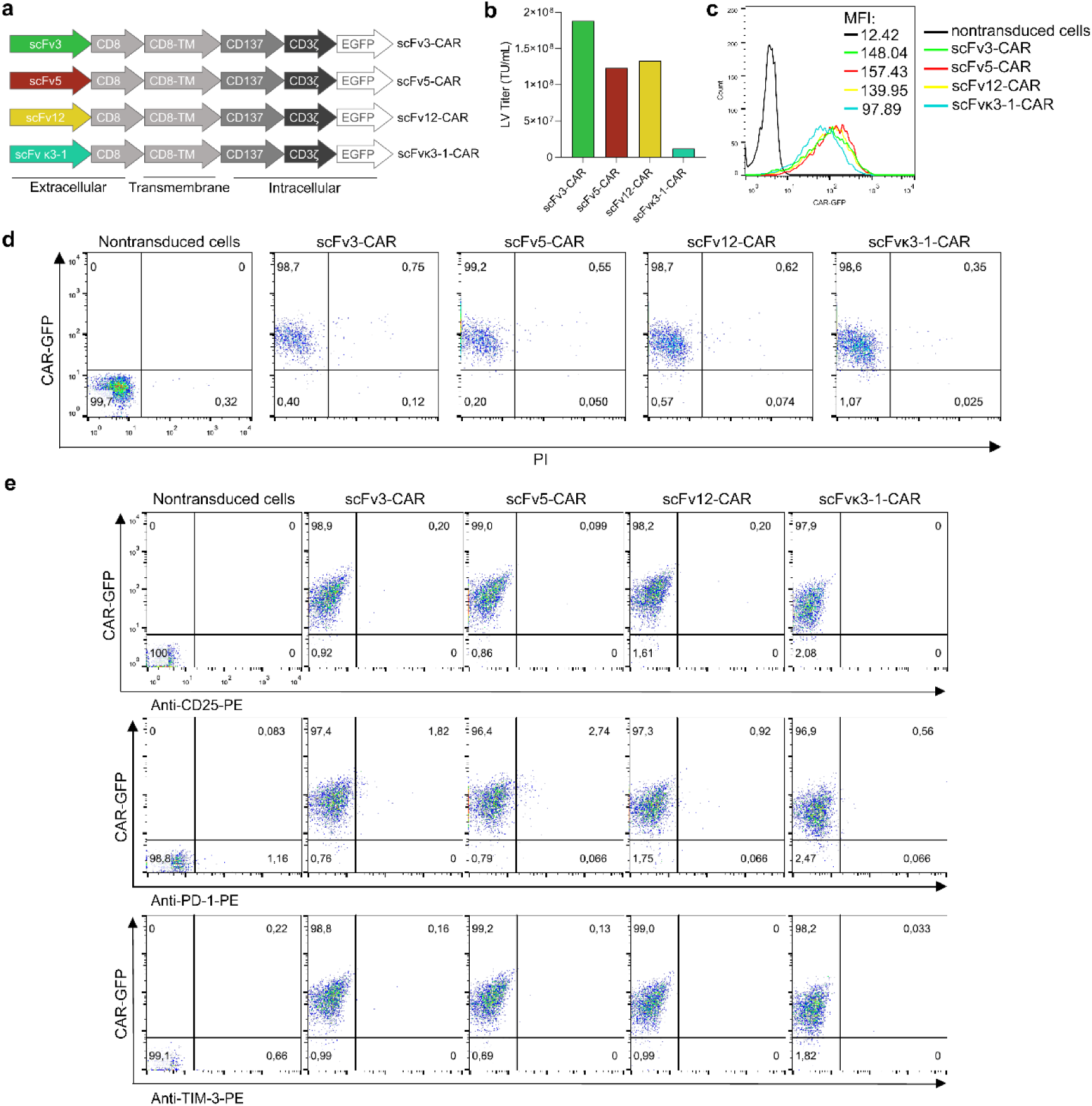
scFv3-CAR, scFv5-CAR, scFv12-CAR, and scFvκ3-1-CAR are highly expressed by Jurkat cells without the induction of an exhausted phenotype. (**a**) Schematic of the domains of the scFv3-CAR, scFv5-CAR, scFv12-CAR, and scFvκ3-1-CAR constructs. (**b**) Quantification of the titer of lentiviral particles containing the plasmid encoding each CAR construct. Jurkat cells were used to determine the titer of the lentiviral particles produced, expressed in transducing units per mL (TU/mL). (**c**) Jurkat cells were modified with an MOI of 5 for scFv3-CAR, scFv12-CAR, and scFvκ3-1-CAR and an MOI of 10 for scFv5-CAR, and the histogram represents the mean fluorescence intensity (MFI) of each CAR construct. (**d**) Modified Jurkat cells were isolated by fluorescence-activated cell sorting, and the dot plots are representative of the frequency of CAR Jurkat cells (GFP expression) and cell viability determined by propidium iodide (PI) staining. **(e)** The cells expressing CAR were incubated with an anti-CD25 monoclonal antibody (PE-conjugated) to evaluate activation and an anti-PD-1 or anti-TIM3 monoclonal antibody (PE-conjugated) to evaluate exhaustion. The percentage of GFP-positive CAR Jurkat cells was determined by flow cytometry, and the dot plots are representative of the results. (**c, d, e**) Nontransduced Jurkat cells were utilized as a negative control.

Jurkat cells were utilized as an initial platform for CAR studies to thoroughly characterize the modified cells, as this model allows for the investigation of various aspects of T cell activation, target recognition, cell exhaustion, and other key processes. Therefore, Jurkat cells were modified to express scFv3-CAR, scFv5-CAR, scFv12-CAR, or scFvκ3-1-CAR, and cellular expansion was performed for 7 days prior to the enrichment of *C. albicans*-specific CAR-T cells for each construct by cell sorting. The expression level of the CAR constructs (Fig. 1c) and their transduction efficiency (Fig. 1d) were assessed through flow cytometry by measuring the mean fluorescence intensity (MFI) and the percentage of viable cells expressing the CAR, respectively. The percentage of Jurkat cells successfully expressing scFv3-CAR, scFv5-CAR, scFv12-CAR, or scFvκ3-1-CAR was at least 96% (Fig. 1c, d).

The induction of the exhausted phenotype of Jurkat cells expressing scFv3-CAR, scFv12-CAR, scFv5-CAR, or scFvκ3-1-CAR was evaluated by measuring the expression levels of PD-1 and TIM-3 via flow cytometry. The frequency of CAR-T cells that were positive for TIM-3 or PD-1 did not differ significantly from those nontransduced cells (Fig. 1e). Furthermore, the expression of the activation marker CD25, which is not constitutively expressed in Jurkat cells, was quantified in *C. albicans*-specific CAR-T cells, and these cells did not exhibit increased CD25 expression (Fig. 1e). These findings demonstrated that Jurkat T cells expressed high levels of scFv3-CAR, scFv5-CAR, scFv12-CAR, or scFvκ3-1-CAR and did not express exhaustion markers, and these modified Jurkat cells could be used to explore the functionality of *C. albicans*-specific CAR constructs against the target.

### scFv5-CAR and scFvκ3-1-CAR mediated the greatest increase in cell activation in response to *C. albicans* exposure

The activation of Jurkat cells expressing scFv3-CAR, scFv5-CAR, scFv12-CAR, or scFvκ3-1-CAR in response to *C. albicans* or soluble antigens was evaluated. *C. albicans*-specific CAR-T cells were cocultured with heat-killed (HK) yeast or hyphal forms of *C. albicans* (at a ratio of 1:1 target to effector cells) and were also incubated with cell wall extracts of *C. albicans*, mannan (α-1,6 backbone with α-1,2- and α-1,3-attached mannose side chains), beta-glucan peptide (BGP), or curdlan. After a 24-hour incubation period, the cell culture supernatants were collected to measure the IL-2 levels by ELISA. Jurkat cells transduced with an empty vector containing only GFP (referred to as Mock cells) served as the negative control (Fig. 2a), while a positive control (PMA + ionomycin or PHA-L + PMA) was used to induce T-cell activation.

**Figure 2.**
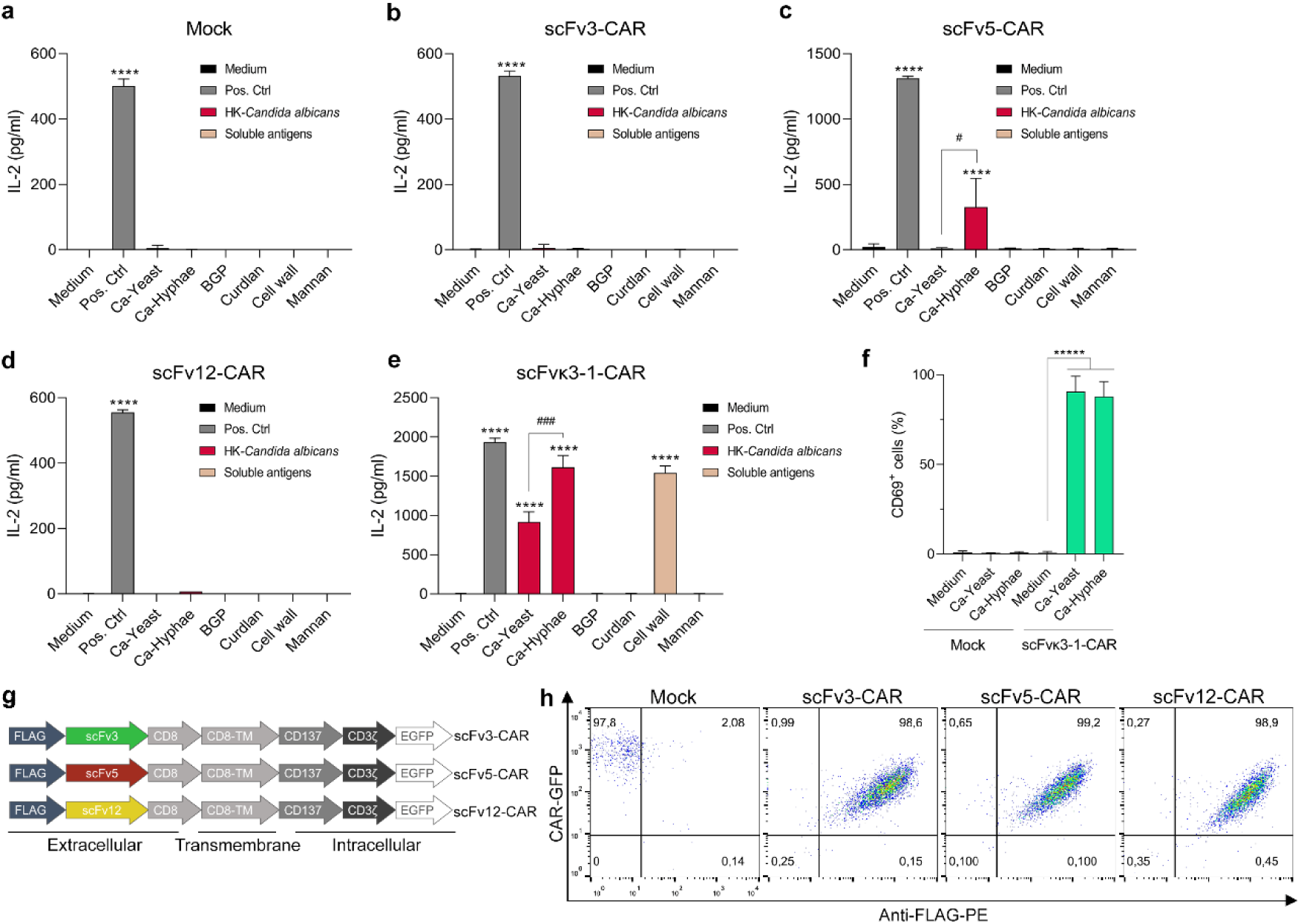
scFv5-CAR and scFvκ3-1-CAR induced the activation of modified Jurkat cells in the presence of *C. albicans.* (**a-f**) The activation assay was performed with Jurkat cells (2 × 10^5^ cells/mL) expressing Mock (**a**), scFv3-CAR (**b**), scFv5-CAR (**c**), scFv12-CAR (**d**), or scFvκ3-1-CAR (**e**). The modified cells were seeded in 96-well plates and cocultured with heat-killed (HK) yeast and hyphae of *C. albicans* (1:1 ratio of target to effector cells) or incubated with mannan (1 µg/mL), curdlan (1 µg/mL), beta-glucan peptide (BGP) (1 µg/mL), or a cell wall extract of *C. albicans* (1 µg/mL). PMA (50 ng/mL) + ionomycin (1 µM) or PHA-L (5 µg/mL) + PMA (50 ng/mL) were used as positive controls to induce cell activation. (**a-e**) After 24 hours of incubation, the cell supernatant was harvested, and the levels of IL-2 were measured by ELISA (*n* = 3–6). (**f**) scFvκ3-1-CAR Jurkat cells and mock control cells were collected after 24 hours of incubation with heat-killed yeast or hyphae of *C. albicans,* and the percentage of CD69-positive cells was detected by flow cytometry (*n* = 3). (**g**) Schematic of the domains of the scFv3-CAR, scFv5-CAR, and scFv12-CAR constructs containing a FLAG tag in the N-terminus of the CAR. (**h**) Dot plots of the detection of the FLAG tag using a PE-conjugated anti-FLAG monoclonal antibody to determine the percentages of modified cells expressing CAR (GFP positive) on the cell surface. One representative experiment out of 3 is presented. In (**a** to **f**), the significance was tested using one-way ANOVA with Dunnett’s test to compare each group with “medium” (*) and Student’s t-test (#). The values are expressed as the mean ± standard deviation (SD). **** p < 0.0001, # p < 0.05, ### p < 0.001

Among all the *C. albicans*-specific CAR constructs evaluated, the scFv5-CAR and scFvκ3-1-CAR induced the highest levels of IL-2 in the presence of *C. albicans* hyphae compared to those induced by the modified cells incubated with medium alone (Fig. 2 b-e). Additionally, scFvκ3-1-CAR induced significant amounts of IL-2 against *C. albicans* yeast and the cell wall extract (Fig. 2e). These data were validated by the high levels of CD69 expressed by scFvκ3-1-CAR-T cells after exposure to yeast and hyphae of *C. albicans*, which was not observed in mock control cells cocultured with *C. albicans* for 24 hours of incubation (Fig. 2f). However, T-cell activation against *C. albicans* or soluble antigens was not induced by the scFv3-CAR or scFv12-CAR (Fig. 2b, d), whereas the scFv5-CAR showed less activation compared to the scFvκ3-1-CAR. In addition, the efficient surface expression of scFv3-CAR, scFv5-CAR, and scFv12-CAR was confirmed in over 98% of the modified cells, as indicated by the detection of the FLAG tag at the N-terminus of the CAR using flow cytometry (Fig. 2g, h), suggesting that the lower activation compared to scFvκ3-1-CAR was not due to insufficient CAR expression.

### *C. tropicalis, C. glabrata,* and *C. auris* were also recognized by scFvκ3-1-CAR

Considering the reduced ability of Jurkat cells expressing scFv3-CAR or scFv12-CAR to recognize both yeasts and hyphae of *C. albicans*, resulting in weak cell activation, subsequent assays focused on CAR constructs containing scFv5 or scFvκ3-1. The growth of Jurkat cells expressing scFv5-CAR or scFvκ3-1-CAR was monitored over time, and the cell concentration and CAR expression were determined every 2 days during 10 days. The expansion rate and percentage of GFP-positive cells did not differ between *C. albicans*-specific CAR-T cells and mock-treated T cells (Fig. 3a, c). However, Jurkat cells showed reduced expression of scFv5-CAR after seven days of culture (Fig. 3b).

**Figure 3.**
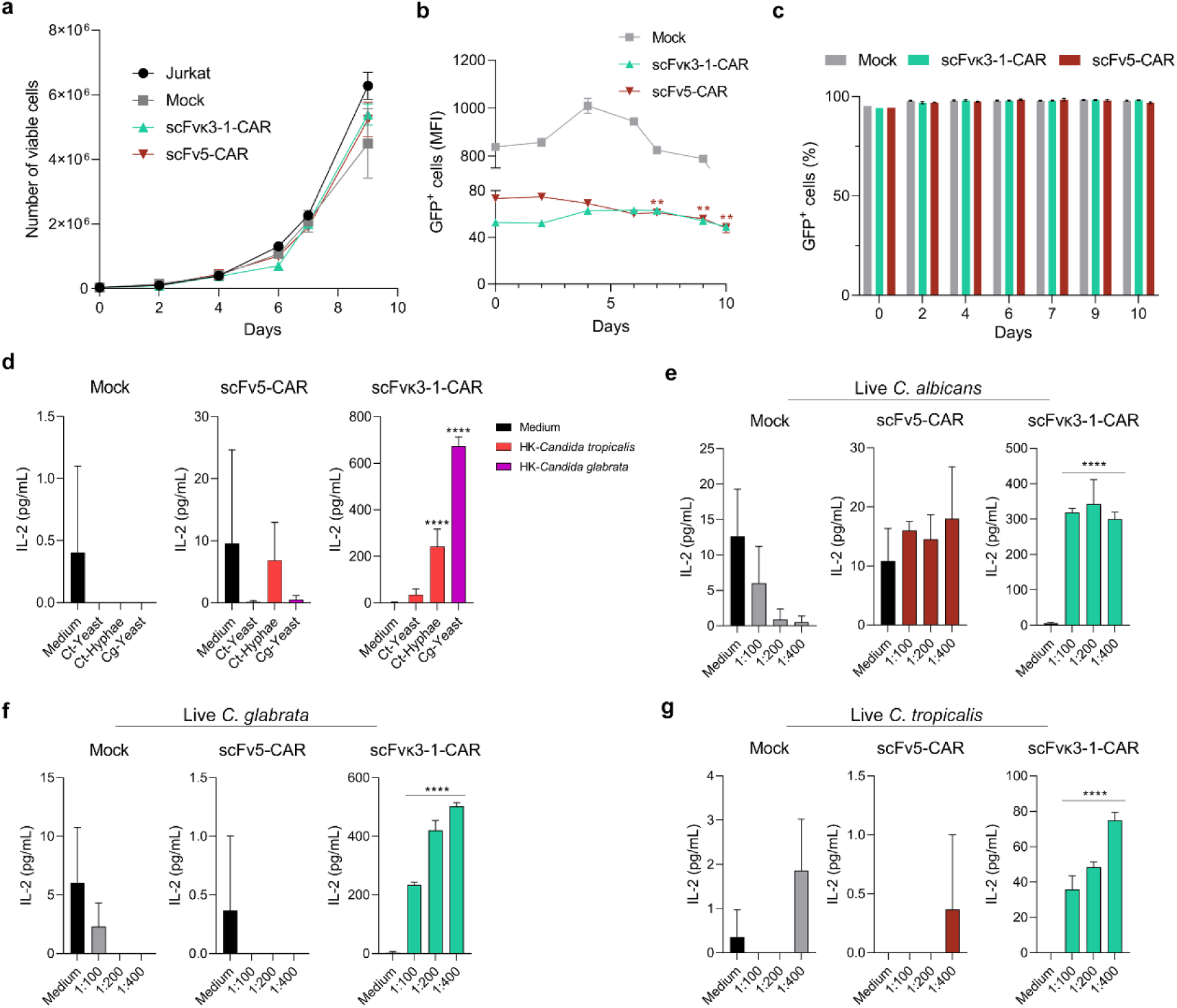
scFvκ3-1-CAR Jurkat cells were activated in the presence of inactivated and live *Candida* species. Jurkat cells expressing Mock, scFv5-CAR, or scFvκ3-1-CAR were cultured for 10 days to determine (**a**) the extent of cellular expansion, (**b**) the level of CAR expression, as determined by the mean fluorescence intensity (MFI), and (**c**) the percentage of GFP-positive cells. (**a-c**) *n* = 3–6 independent experiments. (**d**) Jurkat cells (2 × 10^5^ cells/mL) expressing Mock, scFv5-CAR, or scFvκ3-1-CAR were distributed in a 96-well plate and cocultured with heat-killed (HK) yeast or hyphae of *Candida* spp. (1:1 ratio of target to effector cells). (**e-g**) Modified cells were cocultured with live *Candida* spp. yeasts at different ratios of target to effector cells (1:100, 1:200, and 1:400). (**d-g**) After 24 hours of incubation, the cell supernatants were collected, and IL-2 levels were measured by ELISA (*n* = 3). The groups of each assay were compared to “Medium”. The values are expressed as the mean ± SD. ** p < 0.01, **** p < 0.0001.

The ability of scFv5-CAR and scFvκ3-1-CAR Jurkat cells to recognize different *Candida* species was determined after coculture with heat-killed fungi (at a ratio of 1:1 target to effector cells) for 24 hours. The IL-2 levels in the cell culture supernatant were measured to determine T-cell activation, and scFvκ3-1-CAR induced higher levels of IL-2 in response to yeasts and/or hyphae of *C. glabrata* and *C. tropicalis* than mock cells (Figure 3d). Moreover, scFvκ3-1-CAR-expressing cells also induced cell activation in the presence of clinical isolates of *C. auris* (Fig. S1). In addition, scFvκ3-1-CAR-expressing cells were activated in the presence of live *C. albicans* (Fig. 3e), *C. glabrata* (Fig. 3f), and *C. tropicalis* (Fig. 3g), whereas scFv5-CAR Jurkat cells were not activated in the presence of live *Candida* spp. (Fig. 3e-g). Table S1 lists additional fungal genera, including *Cryptococcus* spp., *Aspergillus* spp., and *Rhizopus* spp., that were cocultured with modified Jurkat cells, but fungal recognition was not detected. Taken together, these data demonstrated that scFv5-CAR recognized only the inactivated hyphae of *C. albicans*, whereas scFvκ3-1-CAR mediated T-cell activation against distinct *Candida* species.

### scFvκ3-1-CAR redirected Jurkat cells to interact with *C. albicans* and induced a strong cell activation

The interaction between scFvκ3-1-CAR Jurkat cells and yeasts or hyphal forms of *C. albicans* was demonstrated through fluorescence microscopy. Heat-killed yeasts and hyphae were stained with Calcofluor white and coincubated with cells modified with scFvκ3-1-CAR or with Mock cells; after 24 hours, the recognition of *C. albicans* by the cells was assessed. The absence of interaction between Mock cells and *C. albicans* was evidenced by the lack of detection of yeasts or hyphae of *C. albicans* in the clusters of Mock cells (Fig. 4). Conversely, scFvκ3-1-CAR Jurkat cells exhibited a strong ability to attract *C. albicans* yeast. The modified cells were in direct contact with the entire length of each hypha structure, showing that both the yeast and hyphae of *C. albicans* were targeted by scFvκ3-1-CAR (Fig. 4).

**Figure 4.**
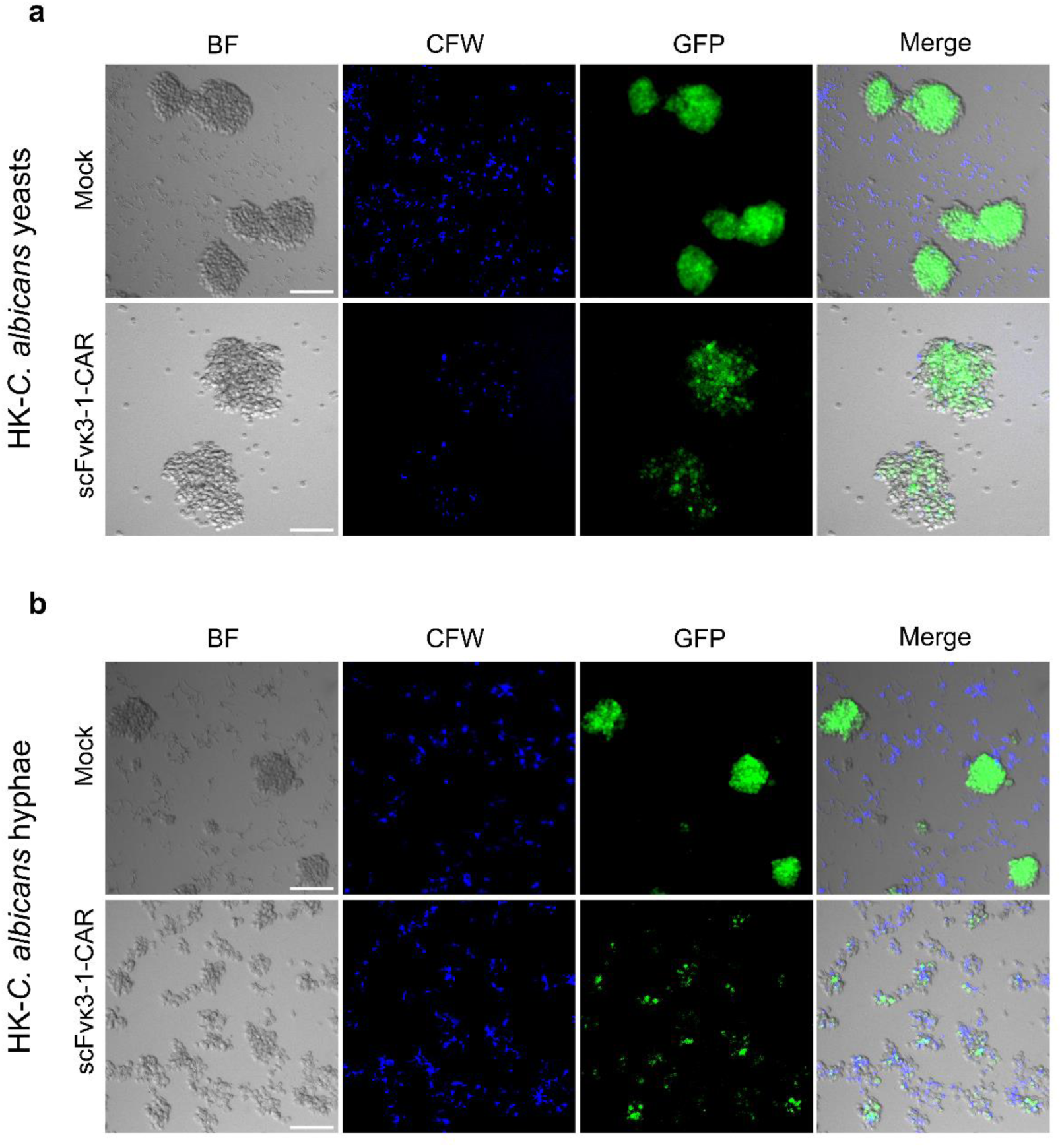
scFvκ3-1-CAR Jurkat cells were redirected to target *C. albicans* yeast and hyphae. Heat-killed (HK) yeasts and hyphae of *C. albicans* (2 × 10^5^ cells/mL) were previously stained with Calcofluor white (CFW) and added to cultures of mock or scFvκ3-1-CAR Jurkat cells at a 1:1 ratio of target to effector cells. After 24 hours, the interaction of Mock cells (GFP^+^) or scFvκ3-1-CAR-modified cells (GFP^+^) with yeast (top panel) or hyphae (bottom panel) of *C. albicans* (blue) was evaluated using fluorescence microscopy (Leica). BF, brightfield. Scale bars, 100 µM.

The intracellular domain of CARs initiates a signaling cascade involving ZAP-70/Syk and Src family of tyrosine protein kinases (TPKs) upon CD3ζ phosphorylation. This signaling cascade was evaluated in scFvκ3-1-CAR Jurkat cells cocultured with heat-killed yeasts or hyphae of *C. albicans* for 10 minutes. Thereafter, the cells were harvested, and phospho-ZAP-70 was detected by flow cytometry using a PE-conjugated monoclonal antibody (Fig. 5a). Hydrogen peroxide (H_2_O_2_) was added to the cell culture as a positive control to induce ZAP-70 phosphorylation. Phospho-ZAP-70^+^ cells were not detected in Mock-Jurkat cells after incubation with *C. albicans*. However, the percentage of phospho-ZAP-70^+^ cells in Jurkat cells expressing scFvκ3-1-CAR increased by 17.13% after coculture with yeast and by 37.59% after coculture with hyphae of *C. albicans* (Fig. 5a).

**Figure 5.**
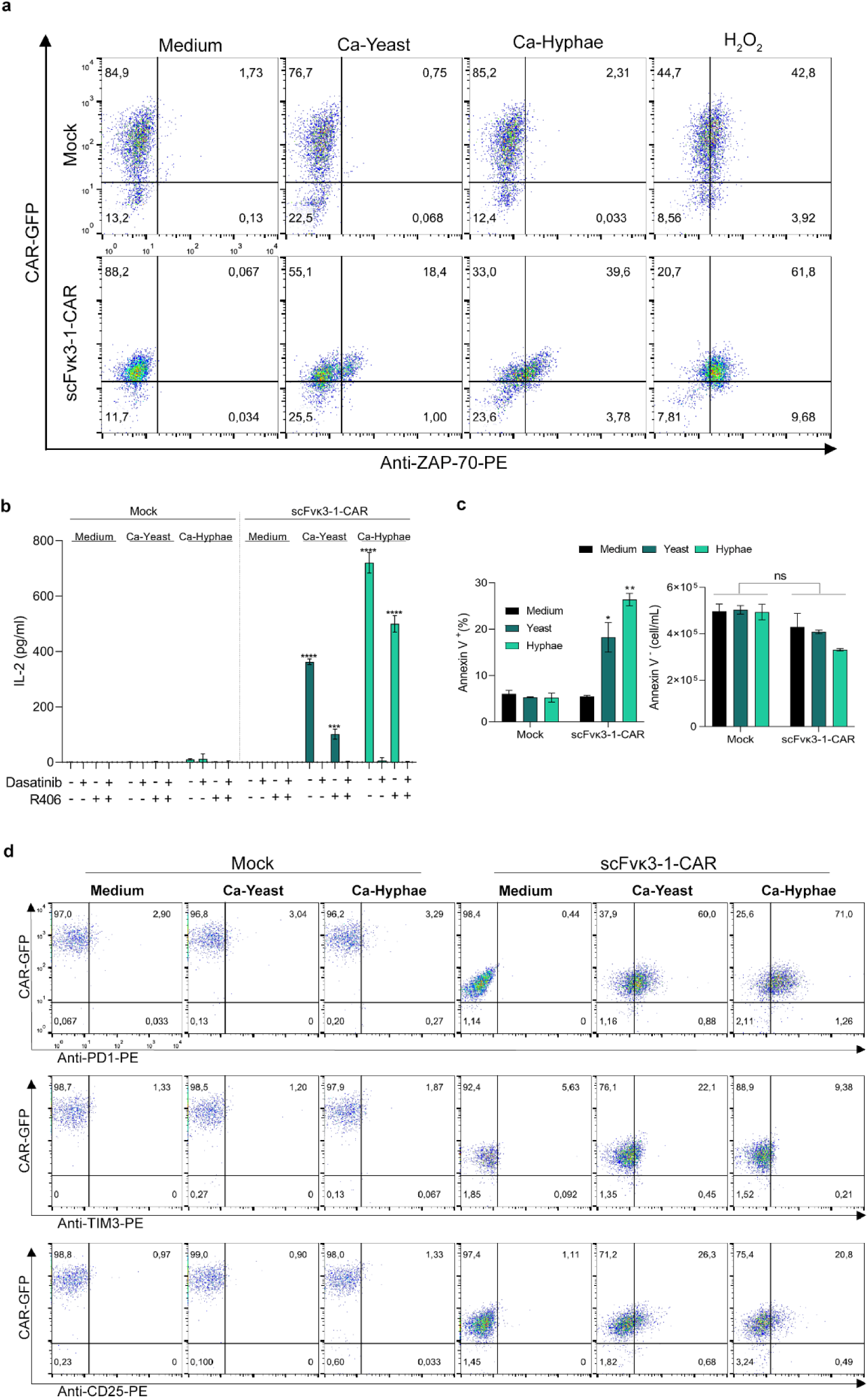
Strong cell activation induced by scFvκ3-1-CAR against *C. albicans* was evidenced by positive staining for phospho-ZAP-70 and exhaustion markers. Jurkat cells (2 × 10^5^ cells/mL) expressing scFvκ3-1-CAR or modified with the Mock construct were cocultured with heat-killed (HK) yeast and hyphae of *C. albicans* in a 96-well plate at a 1:1 ratio of target to effector cells. (**a**) After 10 minutes of incubation, the cells were harvested, fixed/permeabilized, and incubated with an anti-phospho-ZAP-70-PE antibody to determine the percentage of positive cells by flow cytometry. Hydrogen peroxide (0.03%) was used as a positive control for phospho-ZAP-70. One representative experiment out of 2 is presented. (**b**) Modified cells were treated with dasatinib and/or R406 for 3 hours before coculture with *C. albicans*. After 24 hours of incubation, the levels of IL-2 in the cell supernatant were measured by ELISA. (*n* = 3–4). (**c**) Percentage of modified cells positive for Annexin V (left side) and concentration of unstained cells (Annexin V^-^; right side) after 24 hours of coculture with heat-killed yeast and hyphae of *C. albicans* (*n* = 2). (**d**) scFvκ3-1-CAR and mock Jurkat cells were cocultured with distinct forms of *C. albicans* for 24 hours, followed by incubation with PE-conjugated anti-CD25, anti-PD-1, or anti-TIM3 monoclonal antibodies, and the percentages of cells double-positive for GFP and activation or exhaustion markers were determined by flow cytometry. One representative experiment out of 2 is presented. (**b, c**) The experimental groups were compared to the “medium” group in each assay. The values are expressed as the mean ± SD. * p < 0.05, ** p < 0.01, *** p < 0.001, **** p < 0.0001. ns, not significant.

Additionally, Jurkat cells expressing scFvκ3-1-CAR and Mock cells were pretreated with dasatinib (an Src family kinase inhibitor) and/or R406 (a Syk inhibitor) for 3 hours before coculture with *C. albicans*. After 24 hours of incubation, the cell culture supernatants were collected to measure the IL-2 levels, which were significantly increased in cells modified with scFvκ3-1-CAR and cocultured with *C. albicans* yeast and hyphae (Fig. 5b) compared to those in Mock cells. Treatment of scFvκ3-1-CAR Jurkat cells with dasatinib before coculture resulted in complete inhibition of IL-2 production; moreover, treatment with the R406 inhibitor significantly reduced the release of IL-2 by scFvκ3-1-CAR-T cells (Fig. 5b). The combination of both dasatinib and R406 inhibited IL-2 production by cells modified with scFvκ3-1-CAR and incubated with the ligands (Fig. 5b). Taken together, these findings indicated that scFvκ3-1-CAR triggers strong cell activation via the ZAP-70/Syk and Src family kinases.

To determine whether the activated scFvκ3-1-CAR Jurkat cells persist or become exhausted after coculture with *C. albicans*, the cells were analyzed by flow cytometry after staining with Annexin V (Fig. 5c) or labeling for CD24, PD-1, and TIM-3 (Fig. 5d). After 24 hours, the percentage of Jurkat cells expressing scFvκ3-1-CAR and positive for Annexin V increased in the presence of yeast and hyphae (Fig. 5c), whereas the percentage of Annexin V-positive Mock Jurkat cells did not increase. The cell concentration of Annexin V-negative cells was not altered (Fig. 5c). In addition, scFvκ3-1-CAR Jurkat cells cocultured with the yeast or hyphae of *C. albicans* had an increased percentage of cells positive for PD-1, TIM-3, and CD25 compared to cells incubated with medium alone (Fig. 5d). Thus, the strength of the signal transduction triggered by scFvκ3-1-CAR can induce the expression of exhaustion markers in response to target exposure.

### The growth of *C. albicans* was compromised in NK-92 cells expressing scFvκ3-1-CAR

T cell activation mediated by scFvκ3-1-CAR was effectively characterized using Jurkat cells, demonstrating robust activation against *Candida* species. Therefore, we then utilized natural killer (NK)-92 cells to assess ability of scFvκ3-1-CAR to mediate cell activation and effector activity. NK-92 cells were selected due to their inherent cytotoxicity (37, 38) and moderate capacity to damage *Candida* species (44). Additionally, NK-92 cells are already being utilized in clinical trials for solid tumor treatment (39), making this approach promising for future clinical translation.

To further evaluate the potential of scFvκ3-1-CAR to mediate fungicidal activity against *C. albicans*, the NK-92 cell line was modified with scFvκ3-1-CAR using an MOI of 10, and the transduction efficiency was determined by flow cytometry after 3 days. The transduction efficiency of scFvκ3-1-CAR was 40.2% in NK cells, and the expression reached high levels, as represented by the mean fluorescence intensity (MFI) (Fig. 6a). Thereafter, scFvκ3-1-CAR NK-92 cells were isolated by cell sorting, resulting in a high percentage of 93.8% CAR-expressing cells, as indicated by the MFI (Fig. 6a). The interaction between scFvκ3-1-CAR NK-92 cells and live *C. albicans* yeast cells was visualized by fluorescence microscopy following a 6-hour coincubation. Live *C. albicans* yeast cells were frequently detected in the clusters of scFvκ3-1-CAR NK-92 cells, indicating that *C. albicans* yeast was targeted by CAR, which was not observed in nontransduced NK-92 cells (Fig. 6b). These findings were reproduced in the coculture at T:E ratios of 1:50 and 1:100 (Fig. 6b).

**Figure 6.**
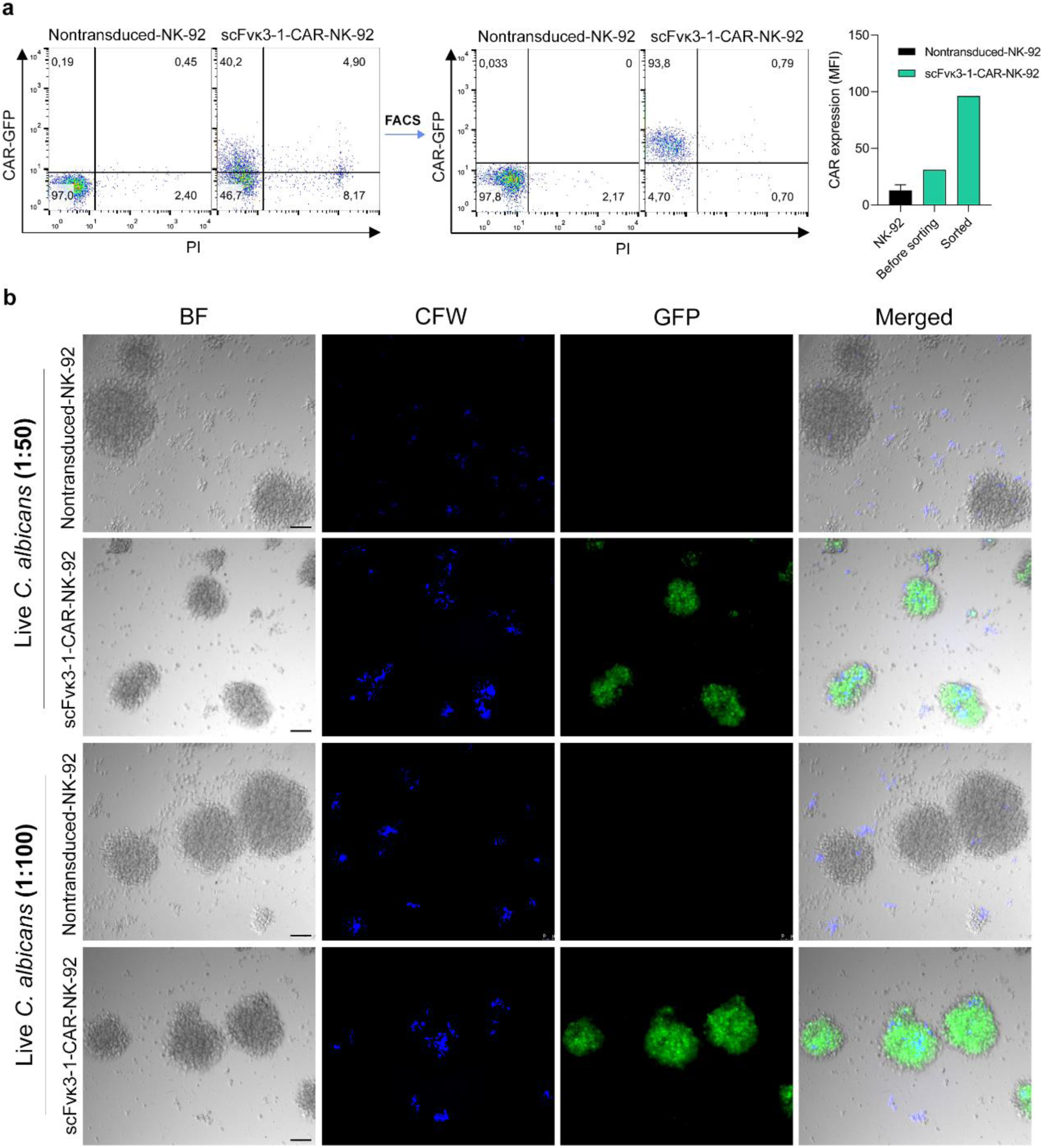
scFvκ3-1-CAR redirected NK-92 cells to interact with live *C. albicans.* (**a**) NK-92 cells were transduced with scFvκ3-1-CAR lentiviral particles at an MOI of 10, and the transduction efficiency was determined by GFP expression using flow cytometry. The modified cells were expanded and enriched by fluorescence-activated cell sorting (FACS). Representative flow cytometry dot plots showing the percentages of scFvκ3-1-CAR-NK cells (GFP^+^). Viable cells were determined by propidium iodide (PI) staining. The bar graphs show the mean intensity fluorescence (MFI) of the scFvκ3-1-CAR-expressing cells (*n* = 1–2). (**b**) NK-92 cells expressing or not expressing scFvκ3-1-CAR (5 × 10^5^/mL) were incubated with live *C. albicans* yeast cells at a ratio of 1:50 or 1:100 (target to effector cells) for 6 hours. Thereafter, Calcofluor white (CFW) was added to the coculture for 15 minutes. The interaction of nontransduced cells (GFP^-^) or scFvκ3-1-CAR-modified cells (GFP^+^) with live *C. albicans* yeast cells (blue) was evaluated using fluorescence microscopy (Leica). BF, brightfield. Scale bars, 100 µM.

To evaluate the activation of modified NK-92 cells, the cells were cocultured with heat-killed yeast and hyphae of *C. albicans* at target-to-effector ratios of 1:50 or 1:100 (Fig. 7a). The IFN-γ levels in the cell culture supernatant were then measured by ELISA after 6 and 24 hours of incubation. Compared with unstimulated cells, unmodified NK-92 cells did not show an increased production of IFN-γ in the presence of heat-killed yeast or hyphae of *C. albicans* (Fig. 7a). In contrast, scFvκ3-1-CAR NK-92 cells produced higher levels of IFN-γ in response to heat-killed yeast or hyphae of *C. albicans* after 24 hours of coculture than scFvκ3-1-CAR NK cells incubated with only medium (Fig. 7a). In addition, NK-92 cells expressing scFvκ3-1-CAR were also activated after coculture with live *C. albicans* at ratios of 1:100 and 1:200 of target to effector cells. This was evidenced by the high levels of IFN-γ released after 24 hours of incubation (Fig. 7b).

**Figure 7.**
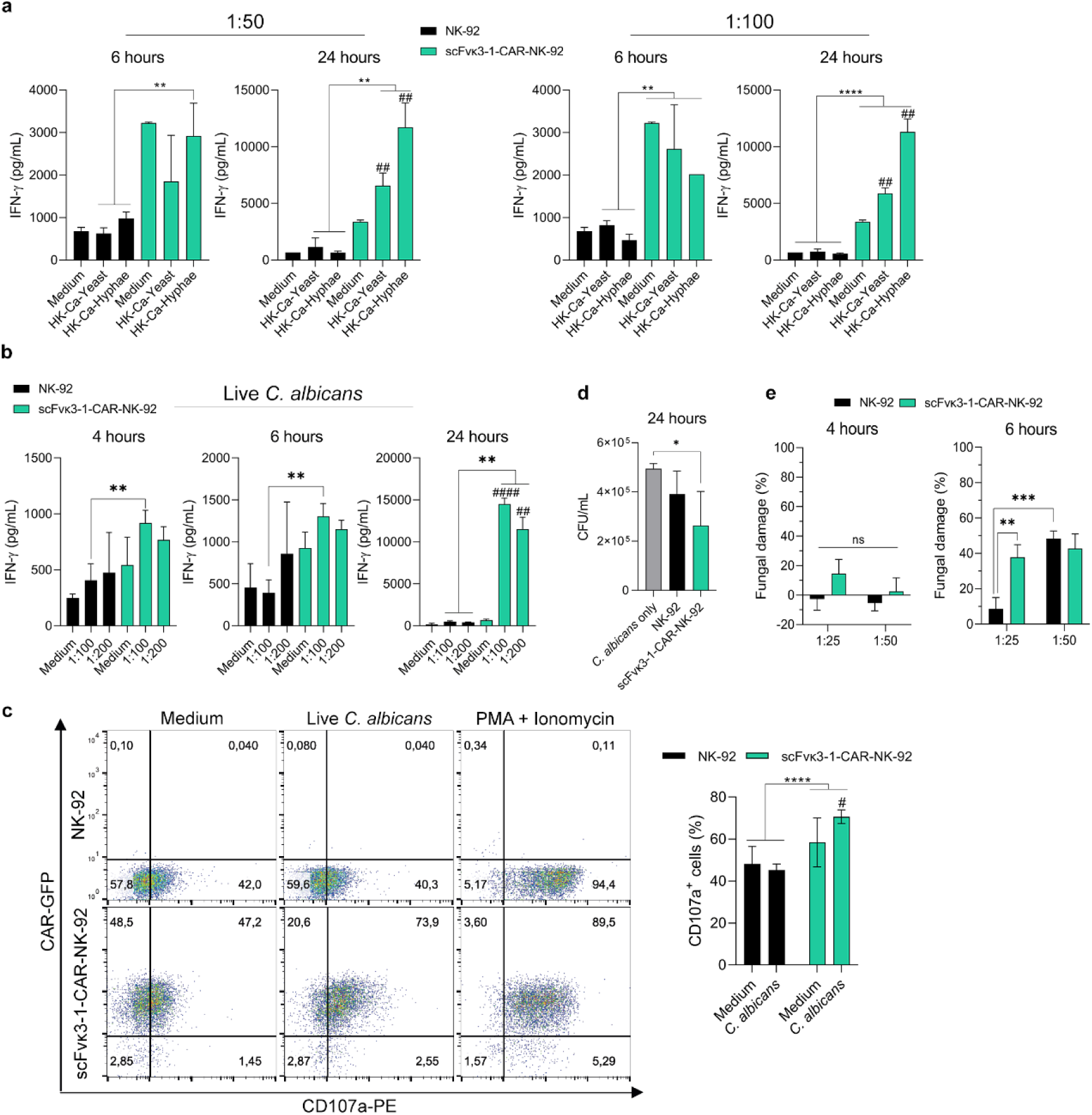
scFvκ3-1-CAR-induced NK-92 cell activation affects *C. albicans* growth. (**a-b**) NK-92 cells expressing or not expressing scFvκ3-1-CAR were cocultured with (**a**) heat-killed (HK) yeast or hyphae of *C. albicans* at a ratio of 1:50 or 1:100 and (**b**) live *C. albicans* yeast cells at a ratio of 1:100 or 1:200 (target to effector cells). After 4, 6, or 24 hours of incubation, the IFN-γ levels (pg/mL) were measured by ELISA. (**a, b**) *n* = 3 independent experiments. (**c**) NK-92 cells modified or not with scFvκ3-1-CAR were cocultured with live *C. albicans* at a ratio of 1:100 (target to effector cells) in the presence of monensin and a PE-conjugated anti-CD107a monoclonal antibody. After 6 hours, the percentage of cells stained for CD107a was determined by flow cytometry. PMA + ionomycin was used as a positive control for degranulation induction, and medium was used as a negative control (*n* = 4). (**d, e**) NK-92 cells modified or not with scFvκ3-1-CAR (5 × 10^5^ cells/mL) were cocultivated with *C. albicans* in a 96-well plate at a ratio of (**d**) 1:100 or (**e**) 1:25 and 1:50. (**d)** After 24 hours, the samples were diluted and plated to perform the colony forming unit (CFU) assay, and the results are presented as CFU/mL. The *C. albicans*-only group refers to the culture of yeast with medium alone (*n* = 3–7). (**e**) After 4 and 6 hours of incubation, NK-92 and scFvκ3-1-CAR-NK-92 cells were lysed with sterile water, and XTT (2,3-bis[2-methoxy-4-nitro-5-sulphenyl]2H-tetrazolium-5-carboxyanilide) solution was added and incubated for 2 hours before the absorbance reading by a spectrophotometer; the results are expressed as fungal damage (%) (*n* = 3). # indicates a significant difference between the experimental group and the “medium” group of scFvκ3-1 CAR-modified cells. The values are expressed as the mean ± SD. #, * p < 0.05; ##, ** p < 0.01; ####, **** p < 0.0001. ns, not significant.

To evaluate the NK cell-mediated cytotoxicity against *C. albicans*, scFvκ3-1-CAR NK-92 cells were cocultured with live *C. albicans* at a 1:100 ratio of target to effector cells, and the release of cytotoxic granules was determined by the cell surface marker CD107a. After 6 hours of incubation in the presence of monensin and an anti-CD107a antibody, the cells were harvested, and the percentage of CD107a^+^ cells was determined by flow cytometry. PMA + ionomycin was used as a positive control to induce strong degranulation. Approximately 40–42% of unmodified NK-92 cells were CD107a^+^ cells in the presence or absence of *C. albicans* (Fig. 7c), whereas the percentage of CD107a^+^ cells among NK-92 cells expressing scFvκ3-1-CAR increased significantly when the cells were incubated with *C. albicans* (73.9%) (Fig. 7c). Taken together, the data demonstrated that scFvκ3-1-CAR mediated strong NK-92 cell activation, suggesting the release of cytotoxic granules and IFN-γ, which are required to combat *C. albicans* infection.

The ability of scFvκ3-1-CAR NK-92 cells to slow or inhibit the growth of *C. albicans* was investigated. Nonmodified NK-92 cells and scFvκ3-1-CAR NK-92 cells were cocultivated with live *C. albicans* yeast at a T:E ratio of 1:100, and *C. albicans* cultivated without NK-92 cells (“*C. albicans* only”) was used as a control (Fig. 7d). After 24 hours of incubation, the samples were plated on Sabouraud dextrose agar media to perform a colony forming unit (CFU) assay. Compared with those in the *C. albicans* only group, the fungal burden in the scFvκ3-1-CAR NK-92 cell group was reduced (Fig. 7d). To evaluate further the fungal damage caused by scFvκ3-1-CAR NK-92 cells after a few hours of *C. albicans* infection, a colorimetric assay was conducted using XTT (2,3-bis[2-methoxy-4-nitro-5-sulphenyl]2H-tetrazolium-5-carboxyanilide). The NK-92 cells expressing or not expressing scFvκ3-1-CAR were cocultured with *C. albicans* at target-to-effector cell ratios of 1:25 and 1:50. After 4 hours of incubation, the level of fungal damage did not significantly differ between the two groups (Fig 7e). However, scFvκ3-1-CAR NK-92 cells exhibited increased fungal damage at a T:E ratio of 1:25 after 6 hours of incubation (Fig. 7e). Additionally, NK-92 cells expressing scFvκ3-1-CAR were incubated with *C. tropicalis* or *C. glabrata,* and after 24 hours of coculture at a T:E ratio of 1:100, the fungal burden in the two groups was unaffected (Fig. S2A, B). These data suggest that scFvκ3-1-CAR NK-92 cells exhibit antifungal activity against *C. albicans*.

### Treatment with scFvκ3-1-CAR NK-92 cells reduced the fungal burden in the kidneys of NSG mice infected with *C. albicans*

NSG mice (NOD scid gamma mice) are commonly used as preclinical models to investigate the efficacy and safety of CAR-T and CAR-NK cell therapy, as these mice lack murine T, B, NK, and functional dendritic cells (45). Initially, *C. albicans* infection was induced in male and female NSG mice to determine the establishment and kinetics of *C. albicans* infection in distinct tissues. For this, the NSG mice were infected with 1 × 10^4^ *C. albicans* yeasts via the retro-orbital route, and the fungal burdens in the liver, kidney, lung, and spleen were measured by CFU assays after 6, 24, and 72 hours of infection (Fig. 8a). The results revealed that *C. albicans* infection was first established in the lung tissue after 6 hours (Fig. 8b). Over the subsequent 72 hours, the infection disseminated, and the greatest fungal burden was observed in the kidneys (Fig. 8b). The fungal burden in the liver remained consistently low throughout all stages of infection, and the fungal burden in the spleen was not affected (Fig. 8b). Interestingly, there was no difference between female and male NSG mice in the establishment and dissemination of *C. albicans* infection over 72 hours (Fig. 8b).

**Figure 8.**
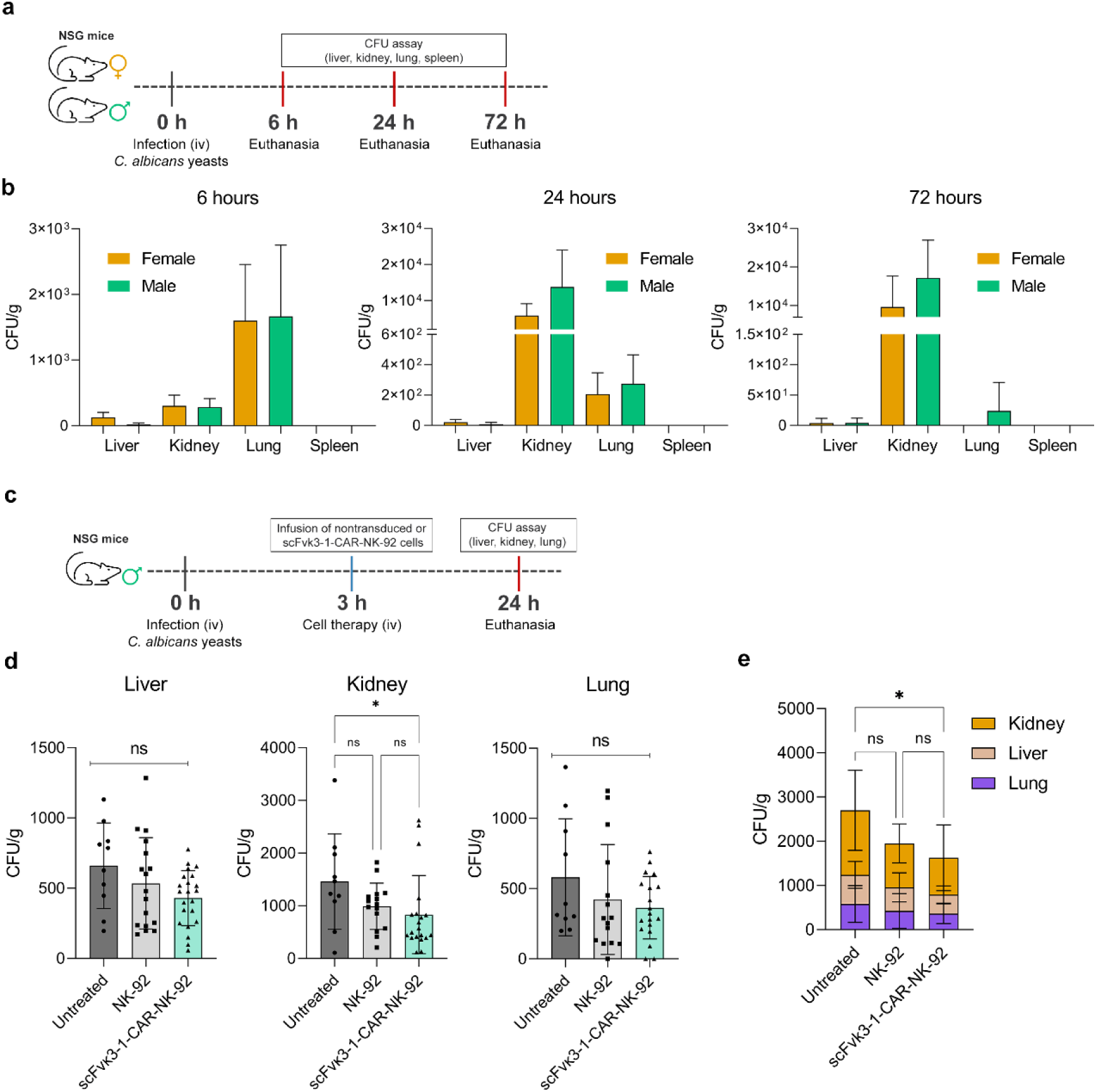
NK-92 cells expressing scFvκ3-1-CAR were redirected to target *C. albicans* in the kidneys. (**a**) Scheme of the *in vivo* biodistribution study of *C. albicans* in NSG mice. (**a,b**) Female and male NSG mice were infected with 1 × 10^4^ *C. albicans* yeast cells via the intravenous route. The mice were euthanized at 6, 24, and 72 hours postinfection to determine the fungal burden in the liver, kidney, lungs, and spleen by CFU assay. The results were normalized relative to the organ mass (g) and are expressed in CFU/g (*n* = 9–10). (**c**) Scheme of the *in vivo* cell therapy for NSG mice infected with *C. albicans.* (**c, d, e)** Male NSG mice were infected with 1 × 10^4^ *C. albicans* yeast cells via the intravenous route, and infusion treatment occurred 3 hours after infection. The NSG mice received NK-92 cells modified or not with scFvκ3-1-CAR (5 × 10^6^ cells/animal) via the lateral tail vein, whereas the untreated mice received only the vehicle PBS. Mice were euthanized at 24 hours postinfection, and their lungs, livers, and kidneys were collected and homogenized to determine the fungal burden by CFU assay (*n* = 10–22). The values are expressed as the mean ± SD. * p < 0.05. ns, not significant.

To evaluate the antifungal efficacy of the modified cells, NSG mice were infected with 1 × 10^4^ *C. albicans* yeast cells via the retro-orbital route, and after 3 hours of *C. albicans* infection, scFvκ3-1-CAR NK-92 cells were intravenously infused via the lateral tail vein (Fig. 8c). Untreated mice were infected with *C. albicans* and received only the vehicle PBS via the lateral tail vein, and another group received unmodified NK-92 cells after *C. albicans* infection (Fig. 8c). Given that NSG mice are immunodeficient, the intravenous infection can rapidly disseminate to multiple organs as showed in Figure 8b, even with the low inoculum used. Therefore, it was necessary to euthanize the mice 24 hours postinfection to prevent them from becoming moribund. Following euthanasia, the lungs, liver, and kidneys were collected and homogenized for CFU assays (Fig. 8c). The results showed a significant reduction in the fungal burden in the kidneys of the mice treated with scFvκ3-1-CAR NK-92 cells compared to that in the kidneys of the untreated mice (Fig. 8d). Although the fungal burden in the liver and lungs did not significantly differ among the groups, the total fungal burden in all tissues combined was lower in the group treated with scFvκ3-1-CAR NK-92 cells than in the untreated group (Fig. 8e). As shown in Figure 8b, intravenous administration of *C. albicans* led to a more localized infection in the kidneys at 24 hours postinfection. This could explain the observed decrease in fungal burden in the kidneys, suggesting that the scFvκ3-1-CAR NK-92 cells were redirected to this organ. Taken together, these results suggested that the infusion of NK-92 cells expressing scFvκ3-1-CAR efficiently targeted *C. albicans* in the kidneys of infected NSG mice, providing proof-of-concept for the ability of scFvκ3-1-CAR to enhance antifungal activity of NK-92 cells.

### scFv κ3-1 has specificity for the mannan of *Candida* spp

The abovementioned findings demonstrated that scFvκ3-1-CAR successfully recognized *C. albicans* yeast and hyphae and induced Jurkat and NK-92 cell activation. scFvκ3-1-CAR incorporates an extracellular domain derived from the scFv region of a human monoclonal antibody fragment (scFv κ3-1) previously generated against *C. albicans* (40). However, the epitope on the *C. albicans* cell wall recognized by scFv κ3-1 is unknown, and carbohydrate was considered to be the possible main target (40). To elucidate this, we generated a scFv κ3-1 protein containing a His-tag in its N-terminus using an expression vector (Fig. S3A). The scFv κ3-1 protein was expressed and purified using a nickel column, followed by size exclusion chromatography (Fig. S3B-C). The protein was then subjected to nuclear magnetic resonance (NMR) analysis to corroborate its proper folding (Fig. S4). The purified scFv κ3-1 protein was used in glycan microarray experiments to determine the monosaccharide or polysaccharide ligands.

The binding of the scFv κ3-1 protein was analyzed using a glycan screening microarray containing saccharide probes, mainly polysaccharides and glycoproteins extracted from fungi and bacteria, among them mannan isolated from *S. cerevisiae* and manno-glycoproteins isolated from *C. albicans* and *A. fumigatus*. The list of the 20 saccharide probes and their sequences are in Table S2. The scFv κ3-1 protein showed exclusively binding to the N-mannoprotein from *C. albicans* (probe #13) characterized by α-1,6-mannose backbone with oligomeric branches of α-1,2-, α-1,3-, and β-1,2-manno-oligosaccharides (46–48) (Fig. 9). This very restricted specificity of the scFv κ3-1 is in sharp contrast with the broader binding profile of the plant lectin Concanavalin A (ConA), a plant lectin with a preference for α-linked mannose, in almost all the mannan-related probes (Fig. 9). scFv κ3-1 protein was further analyzed on a broad-spectrum sequence-defined glycan screening array which contains 672 lipid-linked oligosaccharide probes, including major types of mammalian sequences that occur on glycoproteins (N- and O-linked) and glycolipids carrying various blood group ABO and Lewis antigens together with their sialylated and sulfated analogs, as well as oligosaccharide fragments of glycosaminoglycans and bacterial, fungal and plant polysaccharides (Table S3). The scFv κ3-1 protein exhibited no binding signals with this broad-spectrum screening array (Fig. S5), in contrast to our earlier observation of high-mannose N-glycan binding detected with C-type lectin receptors, including dectin-2 and the macrophage mannose receptor (49). These data highlight the highly specific selective interaction of scFv κ3-1 with the N-mannoprotein of *C. albicans* without reactivity with mammalian-type glycans. Moreover, Haidaris *et al*. showed that scFv κ3-1 did not bind to host skin tissue in model of cutaneous candidiasis, demonstrating its exclusive targeting of *C. albicans* without cross-reactivity (40). Considering that our findings showed that scFvκ3-1-CAR recognized *C. albicans*, *C. tropicalis*, *C. glabrata,* and *C. auris,* it can be inferred that scFv κ3-1 may target the same or similar oligosaccharide moiety in these fungi.

**Figure 9.**
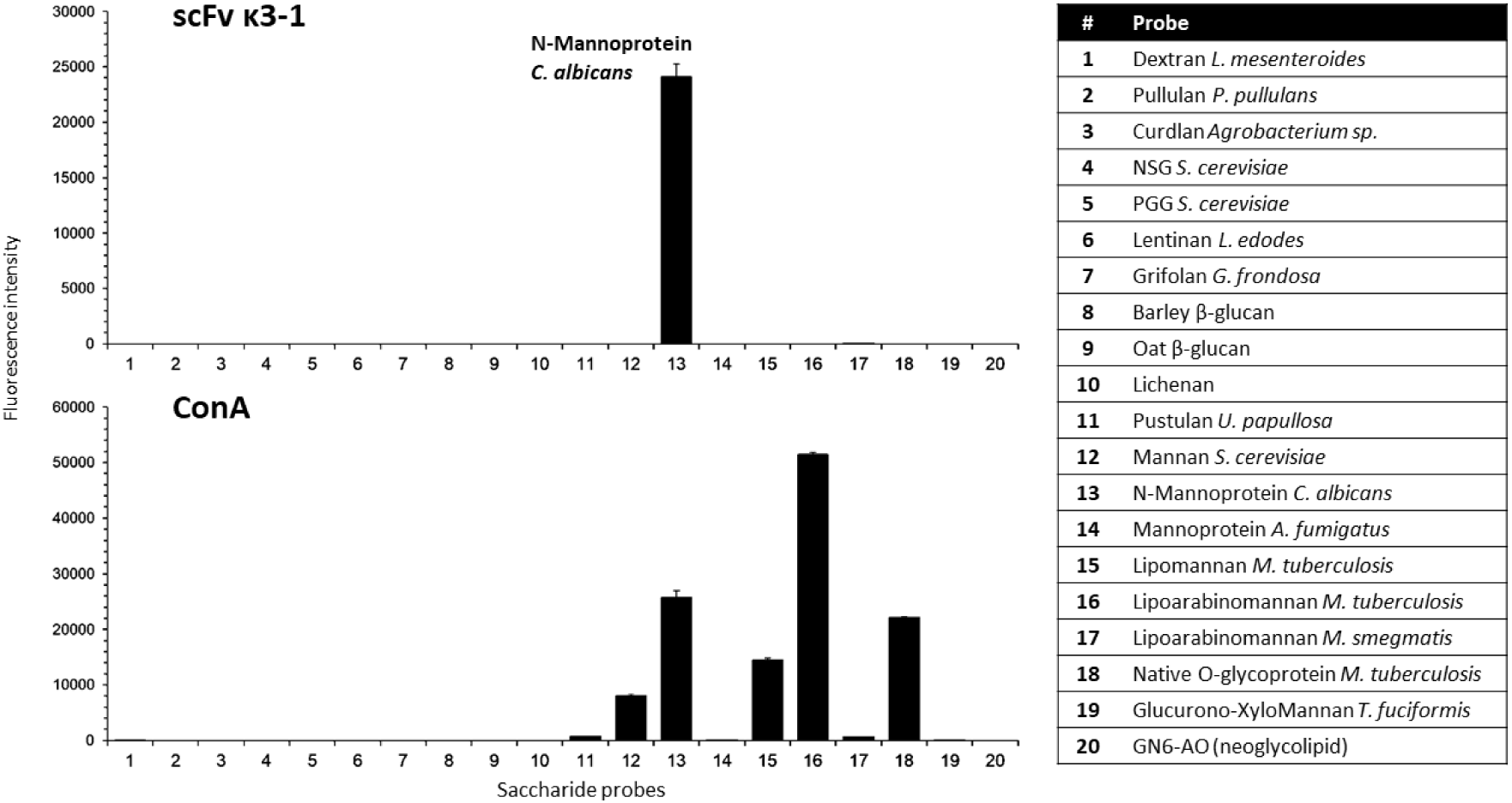
Glycan microarray analysis of scFv κ3-1 and plant lectin ConA using a focused fungal and bacterial polysaccharide array. The 20 saccharide probes are on the right panel, and the information on the oligosaccharide sequences is in Supplementary Table S2. The results are shown at 0.1 ng/spot (saccharides #1–19) and 5 fmol/spot (saccharide #20). The *C. albicans* N-Mannoprotein, which was exclusively bound by the scFv κ3-1 protein (analyzed at 150 µg/mL), is highlighted in the upper panel. Values represent mean fluorescence intensities ± errors (half of the difference of signal intensities of duplicate spots for each saccharide/glycan probe).

## Discussion

Invasive candidiasis, mainly caused by *C. albicans*, poses a major threat to immunocompromised patients, such as those with HIV/AIDS, cancer, or undergoing organ transplantation with prolonged use of immunosuppressive medications. The impacts of invasive candidiasis may be severe, ranging from systemic infection to organ damage, dysfunction, and sepsis. Traditional treatment approaches often require extended hospitalization and the administration of antifungal drugs, which may be ineffective due to their toxicity and the increasing resistance of *Candida* species to these medications (50). The application of CAR technology in cell therapy against IFIs has shown promising results, including a notable reduction in *A. fumigatus* fungal burden *in vivo* and the modulation of virulence factors in *C. neoformans* that are responsible for its evasion of host immunity (29, 30, 32, 33, 36). Here, we demonstrated proof-of-concept for a cell therapy using CAR technology to combat invasive candidiasis. We developed four CAR constructs, scFv3-CAR, scFv5-CAR, scFv12-CAR, and scFvκ3-1-CAR, and expressed these constructs in Jurkat T cells and NK-92 cells. Our results demonstrated that Jurkat cells expressing scFv5-CAR and scFvκ3-1-CAR recognized and induced strong activation in response to *C. albicans*. scFvκ3-1-CAR also recognized several clinically relevant *Candida* species, including *C. tropicalis, C. glabrata*, and clinical isolates of the emerging pathogen *C. auris.* The NK-92 cells expressing scFvκ3-1-CAR were effectively activated when exposed to *C. albicans*, resulting in notable degranulation and reduced fungal viability. Furthermore, infusion of NK-92 cells expressing scFvκ3-1-CAR into NSG mice infected with *C. albicans* reduced the fungal burden in kidneys. Moreover, we found that the mannan of *C. albicans* is the specific antigen recognized by scFvκ3-1-CAR without any cross-reactivity with other glycans.

In this study, we used Jurkat cells, an immortalized human leukemic T-cell line widely used as a platform for CAR studies (51, 52). Our group previously utilized Jurkat cells to analyze various aspects of CAR-T-cell activation, including the expression of activation markers on the cell surface, cytokine production, tonic signaling, and the expression of different CAR constructs (28–31). Therefore, we bioengineered Jurkat cells to express scFv3-CAR, scFv5-CAR, scFv12-CAR, or scFvκ3-1-CAR on the cell surface. The scFv3-CAR and scFv12-CAR constructs did not mediate Jurkat cell activation against either yeast or hyphal forms of *C. albicans,* whereas previous studies had reported that scFv3 and scFv12 recognized *C. albicans*. The low efficiency of cellular activation attributed to scFv3-CAR and scFv12-CAR may be associated with the low stability and low ligand-affinity of the scFv, as previously reported (53, 54). Moreover, scFv3-CAR, scFv5-CAR, scFv12-CAR, and scFvκ3-1-CAR shared identical hinge/transmembrane and signaling domains, and scFvκ3-1-CAR exhibited the strongest ability to induce T and NK cell activation. This finding underscores the impact of scFv domains on CAR functionality against the target. A strategy for further studies would be to enhance the scFv stabilization in CAR constructs by substituting the complementarity-determining regions (CDRs) with different framework regions (FWRs) (55, 56).

The scFv5-CAR exclusively mediated Jurkat-cell activation against *C. albicans* hyphae. *C. albicans*, as a dimorphic fungus, expresses different surface molecules during the transition from yeast to hyphae, which is a critical process for tissue invasion. Therefore, we considered scFvκ3-1-CAR to be more efficient than scFv5-CAR since it recognized both the yeast and hyphae of *C. albicans*, resulting in stronger cell activation. Immune cells expressing scFvκ3-1-CAR may inhibit *C. albicans* switching, thereby increasing host survival (57, 58). Further experiments should be performed to investigate whether scFvκ3-1-CAR-expressing modified cells can effectively suppress this yeast-to-hypha transition. Another important finding was the activation of scFvκ3-1-CAR Jurkat cells in the presence of other *Candida* species, highlighting their broad recognition capability. Moreover, scFv κ3-1 was reported to recognize *Candida dubliniensis* (40), which was not included in our activation assays.

High levels of IL-2 in the culture supernatant and upregulation of CD69 and CD25 were observed when Jurkat cells expressing scFvκ3-1-CAR were cocultured with *C. albicans* yeast and hyphae, indicating full T-cell activation. Additionally, the activation of cells was validated through analysis of exhaustion markers, which offer valuable insights into the functional status of engineered cells and are commonly observed as recurring indicators of cellular activation in response to target exposure and binding. Accordingly, the upregulation of PD-1 and TIM-3 was observed following cell activation. However, these markers were not detected in the absence of *C. albicans*, suggesting that the expression of CAR alone did not lead to cell exhaustion. Previous studies have shown that exhausted CAR-T cells are less effective at killing targets and proliferating, and PD-1 and TIM-3 expression is required to attenuate strong stimulation (59, 60). Within 24 hours of incubation with the targets, scFvκ3-1-CAR Jurkat cells actively produced IL-2 and exhibited signs of exhaustion. However, compared with control cells, scFvκ3-1-CAR Jurkat cells did not show a higher apoptosis rate, but prolonged activation could lead to functional exhaustion over time. The enhanced exhaustion markers in CAR-T cells can be attributed to excessive signaling transduction from the CAR. Mutating two out of the three immunoreceptor tyrosine-based activation motifs (ITAMs) on the CD3ζ of CAR may potentially alleviate cell exhaustion in CAR-induced T-cells (61). Importantly, scFvκ3-1-CAR incorporates CD137 in the costimulatory domain, and CAR-T cells incorporating CD137 have been demonstrated to have superior persistence, reduced exhaustion, and enhanced tumor control compared to those incorporating CD28 (62, 63).

The antifungal efficacy mediated by CARs was analyzed by modifying NK-92 cells, a cell line that currently plays a crucial role in CAR studies due to its cytotoxicity against tumor cells (37, 38), (39). Previous studies have used various mouse strains to establish experimental models of invasive candidiasis, and the role of NK cells in protecting these mice can be complex, exhibiting either beneficial or detrimental effects depending on the experimental conditions (64, 65). A recent study revealed that depletion of NK or T cells in C57BL/6 mice infected with *C. albicans* yeast increased mortality and fungal burden in the kidney, indicating that both NK and T cells are important in controlling invasive candidiasis (66). Another study showed that NK cells can contribute to hyperinflammation and can exacerbate the disease in immunocompetent mice (64). Thus, there is a need for further studies on the role of NK cells in the context of invasive candidiasis to elucidate the mechanism involved in antifungal activities mediated by NK cells. In this study, activated NK-92 cells expressing scFvκ3-1-CAR released high levels of IFN-γ in the culture supernatant in response to *C. albicans* yeast and hyphae. The production of IFN-γ by scFvκ3-1-CAR NK-92 cells in the absence of *C. albicans* suggested the presence of tonic signaling. Tonic signaling can be caused by antigen-independent clustering of CARs in T cells, which is associated with a high density of CARs on the cell surface and self-aggregation. Notably, we transduced Jurkat cells with scFvκ3-1-CAR at an MOI of 5, while we transduced NK-92 cells at an MOI of 10, which may have contributed to tonic signaling. A previous study from our group suggested a correlation between the expression level of CAR and the strength of tonic signaling, which increased with increasing MOI (36). Several studies have investigated the benefits of tonic signaling in promoting the expansion of T cells *in vivo* and *in vitro* (63, 67), which we also observed in NK-92 cells expressing scFvκ3-1-CAR *in vitro*. Another important activation marker for scFvκ3-1-CAR-NK-92 cells was the upregulation of CD107a, indicating cell degranulation in the presence of *C. albicans.* CD107a upregulation may correlate with increased perforin levels, which reduce the metabolic activity of *C. albicans* hyphae and inhibit the elongation of *C. albicans* filaments (16, 17). In contrast to observations on primary NK cells (44), unmodified NK-92 cells did not exhibit CD107a upregulation after co-culture with *C. albicans* for 6 hours. Further experiments should be performed to evaluate the levels of perforin, granulysin, and granzyme B in the supernatant of scFvκ3-1-CAR-NK-92 cells cocultured with *C. albicans* to fully characterize the cytotoxic activity of these modified cells. The XTT assay showed that at 4 hours after infection, coincubation with nonmodified NK-92 cells led to an increase in the percentage of metabolically active *C. albicans,* whereas coincubation with modified cells at a 1:25 ratio resulted in approximately 20% fungal damage. At 6 hours after infection, scFvκ3-1-CAR-NK-92 cells caused more fungal damage than nonmodified NK-92 cells.

This study used NSG mice as an experimental model for invasive candidiasis. NSG mice are typically used to investigate the safety and efficacy of CAR-T and CAR-NK cell therapy due to their lack of functional T, B, and NK cells. However, only one previous study has used NSG mice as an experimental model for invasive candidiasis (64). Therefore, we first characterized the kinetics of *C. albicans* infection in NSG mice. A previous study used bioluminescent *C. albicans* reporter strains in BALB/c mice to analyze infection kinetics *in vitro* and *in vivo* (68). The authors observed transient *C. albicans* localization in the lungs at 8 hours postinfection, with an increase in CFU observed in the kidneys by 24 hours (68). The mice were euthanized at 72 hours due to the development of severe disease (68). Our findings in NSG mice align with those results, except that we observed transient *C. albicans* in the lungs at 6 hours postinfection. Evaluating the kinetics of *C. albicans* infection in NSG mice revealed migration from the lungs to kidneys by 6-8 hours postinfection. Then, the scFvκ3-1-CAR-NK-92 cell therapy was initiated after 3 hours of infection to investigate its efficacy in controlling pulmonary fungal burden and limiting dissemination. The current CAR-NK therapy protocol proposed for the treatment of invasive candidiasis was unable to reduce the burden of *C. albicans* in the lungs. However, scFvκ3-1-CAR-NK-92 cells were responsible for reducing fungal burden in the kidneys of mice infected with *C. albicans* compared to those in the untreated group. Previous studies have indicated that there is an immediate NK cell migration to the lungs after intravenous infusion, followed by accumulation of these cells in the kidneys after 4 hours, which is facilitated by C-C chemokine receptor type 5 (CCR5)-mediated recruitment during *C. albicans* infection (69, 70). While NK cells are critical for fungal clearance, their heavy accumulation in kidneys relative to the lungs or liver may have contributed to the observed lack of cell targeting in these two organs. Moreover, the formation of hyphae, which is the main target for scFvκ3-1-CAR-NK-92 cells rather than yeast cells, typically emerges after 24 hours of *C. albicans* infection *in vivo* (71), coinciding with the time point at which increased scFvκ3-1-CAR-NK-92 cell activation was observed *in vitro*. Thus, our *in vivo* and *in vitro* observations indicate that the redirection of NK-92 cells modified with scFvκ3-1-CAR to combat *C. albicans* dissemination to the kidneys occurs after 24 hours of infection. Adjustments in this therapeutic strategy may be needed to enhance fungal burden clearance by scFvκ3-1-CAR-NK-92 cells in different organs, and multiple NK cell infusions, a strategy already utilized in the treatment of solid tumors with CAR-NK-92 cells, may enhance antifungal efficacy (72, 73). Additionally, experiments should be performed to investigate the benefits of combining the infusion of NK-92 cells expressing scFvκ3-1-CAR with antifungal drugs. This association between cell therapy and antifungal drugs may considerably decrease the minimum inhibitory concentration (MIC) of antifungal drugs, such as caspofungin, and reduce the fungal burden systemically without causing severe side effects.

Knowing the scFvκ3-1-CAR-mediated antifungal activity in NK cells, we focused on characterizing the specific antigen of *C. albicans* recognized by the scFvκ3-1-CAR. A previous study that characterized scFv κ3-1 described epitope sensitivity to periodate treatment but not proteinase K treatment, suggesting that scFv κ3-1 recognizes carbohydrates (40). Therefore, we generated the scFv κ3-1 protein without the other CAR domains to evaluate its binding using glycan microarray analysis. Notably, scFv κ3-1 showed high-specific binding to the mannan of *C. albicans,* without any cross-reactivity with other glycans. Haidaris et al. previously reported that scFv κ3-1 specifically bound to *C. albicans* in mice with cutaneous infection (40). There was no binding to mouse skin, corroborating to its specificity (40). The scFv κ3-1 did not bind to the mannans of *S. cerevisiae* or *A. fumigatus,* suggesting that there were structural differences in these mannans. *A. fumigatus* has highly branched mannans covalently bound to a glucan-chitin core (74), whereas *S. cerevisiae* mannan comprises an α-1,6 backbone with α-1,2 and α-1,3-mannose side branches (75). These results are consistent with our observation that scFvκ3-1-CAR-Jurkat cells are not activated in the presence of soluble *S. cerevisiae* mannan. Importantly, scFvκ3-1-CAR binds to *C. albicans*, *C. tropicalis, C. glabrata,* and *C. auris*, suggesting the presence of a conserved epitope among these species. In this context, *C. albicans, C. glabrata,* and *C. tropicalis* share antigenic factor 6 in their mannan structure. This term is used for oligosaccharides containing one to four β-1,2-linked mannose units connected to α-1,2-linked mannotetraose (75, 76). *C. auris* also has oligosaccharides containing α-1,2- and β-1,2-linked mannose units in its mannan structure (77). Moreover, compared to yeast cells, the *C. albicans* hyphal form contains more of the β-1,2-linked mannose residues associated with antigenic factor 6, which may be related to the stronger binding of scFv κ3-1 to hyphae than to yeasts (78). Therefore, it is likely that the antigenic factor 6 from the mannan of *Candida* spp. constitutes the epitope for scFvκ3-1-CAR recognition, associated with the β-1,2-linked mannose residues, leading to the activation events in Jurkat and NK-92 cells expressing scFvκ3-1-CAR, as proposed in Figure 10.

**Figure 10.**
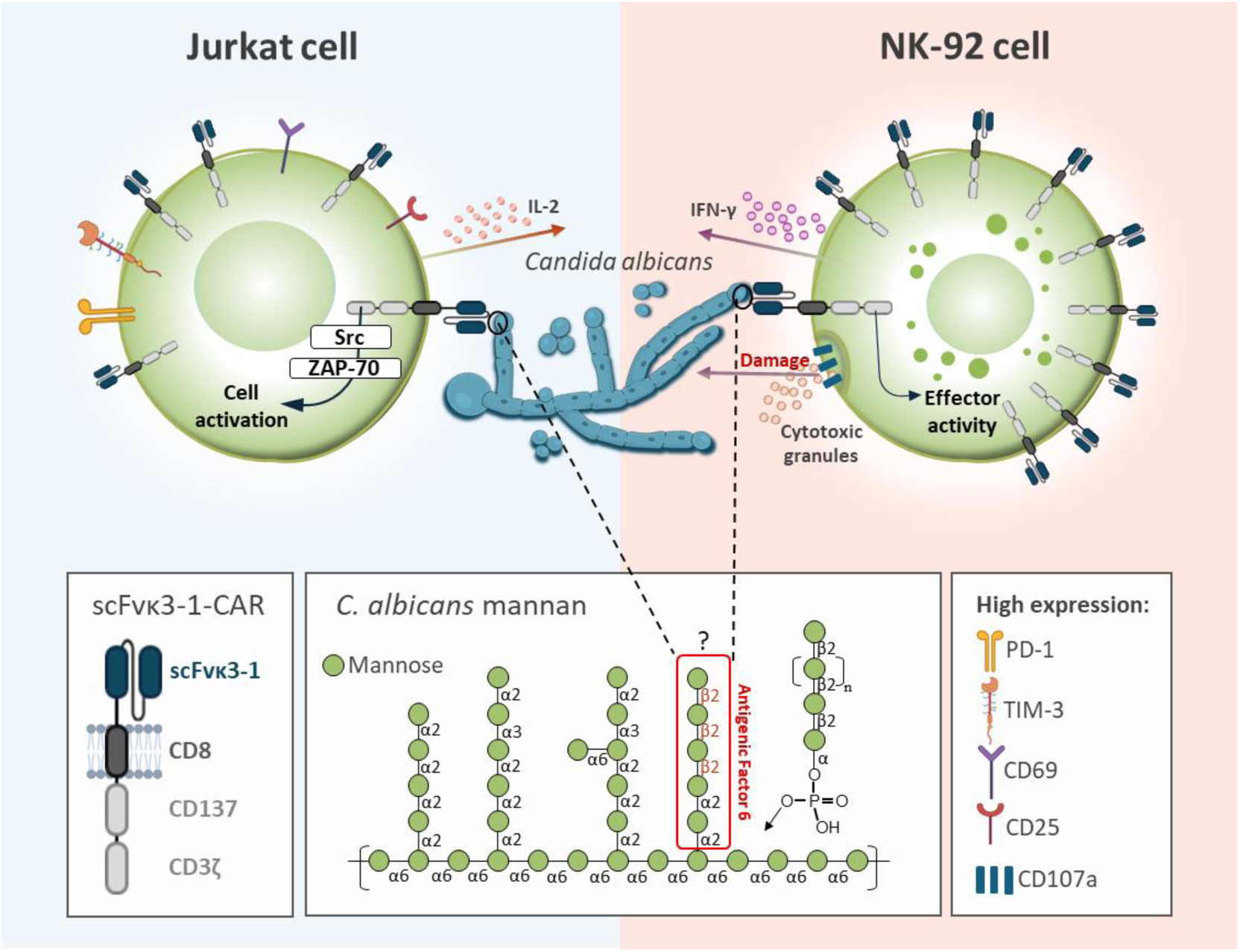
Proposed model for scFvκ3-1-CAR-mediated cell activation upon recognition of *C. albicans.* The structure of scFvκ3-1-CAR comprised CD3ζ and CD137 in the intracellular domain, CD8 in the transmembrane and hinge domains, and scFvκ3-1 in the extracellular domain. Upon target recognition, scFvκ3-1-CAR mediated activation in both Jurkat and NK-92 cells. In Jurkat cells, the scFvκ3-1-CAR activation initiated a signaling cascade involving the phosphorylation of Src family tyrosine protein kinases and ZAP-70/Syk, leading to high IL-2 production and elevated expression of activation markers CD69 and CD25. These events induced an exhaustion phenotype in Jurkat cells, evidenced by the expression of exhaustion markers PD-1 and TIM-3. In NK-92 cells, the scFvκ3-1-CAR activation resulted in high IFN-γ production and elevated expression of CD107a, which is associated with increased release of cytotoxic granules. The effector activity of scFvκ3-1-CAR-NK-92 cells was demonstrated by the increased fungal damage to *C. albicans* and improved clearance in the kidneys of NSG mice. scFvκ3-1-CAR recognizes the mannan of *C. albicans* yeast cells and hyphae, identified by glycan microarray analyses, and antigenic factor 6 (comprising one to four β-1,2-linked mannose units connected to α-1,2-linked mannotetraose (79)) is the potential moiety recognized by scFvκ3-1-CAR.

In conclusion, this study reported a novel CAR that recognizes *C. albicans* yeast and hyphae. scFvκ3-1-CAR mediated Jurkat and NK-92 cell activation in response to *C. albicans* and other *Candida* species, and the cytotoxic activity of scFvκ3-1-CAR NK-92 cells against *C. albicans* was evident *in vitro* and *in vivo*. Overall, the findings demonstrate that scFvκ3-1-CAR is a promising construct for mediating the activation of primary cells targeting *Candida* spp., offering valuable insights for potential future clinical translation in the treatment of invasive candidiasis.

## Materials and Methods

### Chemicals and reagents

Chemicals and reagents for cell culture were obtained from Thermo Fisher Scientific (sodium pyruvate, XTT), Sigma‒Aldrich (penicillin–streptomycin, dasatinib, ionomycin, folic acid, myo-inositol, menadione, curdlan, mannan), Gibco (MEM-alpha, fetal bovine serum, horse serum, PBS, 2-mercaptoethanol), and HyClone (RPMI 1640, high-glucose DMEM). Reagents for the fungal culture experiments were obtained from KASVI (Sabouraud dextrose), Sigma‒Aldrich (Agar, Calcofluor white), and AppliChem (Kanamycin). Reagents and chemicals for flow cytometry, microscopy, and biological activity assays were obtained from BD Biosciences (Fixation/Permeabilization Kit, monensin, cytokine kit, Annexin V cat. 556422, anti-CD25 (cat. 555432), anti-CD69 (cat. 555531), anti-TIM3 (cat. 563422), anti-PD-1 (cat. 560795), anti-phosZAP70 (cat. 557881), anti-CD107 (cat. 555801) antibodies), Sigma‒Aldrich (propidium iodide, anti-FLAG, PE-conjugated streptavidin), and InvivoGen (R406 inhibitor, β-glucan peptide). The transfection reagent and lentiviral particle concentration solution were obtained from Invitrogen (Lipofectamine 3000) and Takara (Lenti-X concentrator), respectively.

### Cell lines and growth conditions

HEK-293T and Jurkat cell lines were obtained from the Rio de Janeiro Cell Bank (BCRJ). The NK-92 cell line was acquired from the American Type Culture Collection (ATCC). HEK-293T cells were cultured in high-glucose DMEM supplemented with fetal bovine serum (10%), sodium pyruvate (100 mM), and penicillin–streptomycin (1%). Jurkat cells were cultured in RPMI 1640 medium supplemented with fetal bovine serum (10%), sodium pyruvate (100 mM), and penicillin–streptomycin (1%). NK-92 cells were cultivated in MEM supplemented with fetal bovine serum (12,5%), horse serum (12,5%), penicillin–streptomycin (1%), 2-mercaptoethanol (55 mM), folic acid (7 μg/mL), myoinositol (36 μg/mL) and IL-2 (200 U/mL). Cells were cultured in a humidified atmosphere with 5% CO_2_ at 37 °C.

### Animals

Nine-week-old female and male NSG mice were maintained in the animal facility situated at the Blood Center of the Ribeirao Preto Medical School of the University of São Paulo, Brazil. To perform the animal studies, the mice were housed at the Department of Cell and Molecular Biology and Pathogenic Bioagents of Ribeirão Preto Medical School, and animal procedures were conducted according to the Ethical Principles Guide for the Care and Use of Laboratory Animals adopted by the Brazilian College of Animal Experimentation. The Committee on Ethics in Animal Research of Ribeirão Preto Medical School approved this study (protocol 071/2021). The animals were housed under controlled environmental conditions at ambient temperature and humidity and maintained on a 12-hour light cycle. The mice were provided ad libitum access to chow and water. The mice were anesthetized by administration of ketamine hydrochloride (20.0 mg/kg, intraperitoneal) and xylazine hydrochloride (2 mg/kg, intraperitoneal) before euthanasia after the *C. albicans* infection period.

### Culture of *Candida* spp. and *Cryptococcus* spp. and heat-killing of yeast and/or hyphal and pseudohyphal forms

*Candida albicans* (ATCC 64548)*, Candida parapsilosis* (ATCC 90018)*, Candida tropicalis* (H2747)*, Candida glabrata* (H3479)*, Candida krusei* (ATCC 6258)*, Candida guilliermondii* (ATCC 22017)*, Candida auris* (clinical isolates 467, 468 and 469)*, Cryptococcus gattii* (R265)*, Cryptococcus neoformans* (H99) and *Cryptococcus neoformans* (acapsular mutant, CAP67) were recovered from 25% v/v glycerol stocks stored at -80 °C. The fungi were spread on Sabouraud dextrose agar, and after 24 hours of incubation at 30 °C, the yeasts were inoculated into Sabouraud dextrose broth and were agitated at 150 rpm at 30 °C for 16 hours. The yeasts were harvested by centrifugation at 7600 × g for 10 min at 25 °C and washed twice with sterile PBS, and the cell concentration was determined using a Neubauer chamber. The yeast-to-hyphal/pseudohyphal form transition was achieved by seeding 2 × 10^5^ yeast cells/mL (4 × 10^4^ yeast cells/well) in 96-well plates containing RPMI-1640 or MEM. The yeast cells were incubated at 37 °C for 3-4 hours, and the transition stage was monitored and confirmed using an inverted microscope. Heat-killing (HK) of hyphal and pseudohyphal was performed by heating 70 °C for 1 hour. The cell culture supernatants were replaced with the fresh medium before adding Jurkat or NK-92 cells modified with or without CAR. *Candida* spp. and *Cryptococcus* spp. yeast forms were inactivated at 70 °C for 1 hour, and the yeast cells were washed twice with sterile PBS before cell counting using a Neubauer chamber.

### CAR construction

The antigen binding domain, hinge/transmembrane domain, and signal transduction regions that compose the CAR were synthesized by GenScript and inserted into a lentiviral plasmid backbone (pLentiCas9-EGFP, GenScript, NJ, USA) using AfeI/BamHI restriction sites, and the GFP reporter gene was inserted into the CAR reading frame. The scFvs in the antigen binding domain were derived from monoclonal antibody fragments previously described (40–42), which generated scFv3-CAR, scFv5-CAR, scFv12-CAR, and scFvκ3-1-CAR constructs. The hinge/transmembrane domain was composed of the CD8 molecule (UniProt P01732, 136–206 aa position) and the cytoplasmic portion of the human CD137 molecule (UniProt Q07011, 214–255 aa position) coupled to the cytoplasmic region of the human CD3ζ molecule (UniProt P20963, 52–164 aa position). The scFv3-CAR, scFv5-CAR, and scFv12-CAR each contained a FLAG tag in the N-terminus. The Mock construct was a plasmid backbone containing the GFP sequence alone and served as a control lentiviral (pLenti-mock) empty vector.

### Production of lentiviral particles

HEK-293T cells (2×10^6^ cells/flask) were dispensed in 25 cm^2^ cell culture flasks to produce lentiviral particles containing the plasmid encoding CAR. The cells were cotransfected with the accessory plasmids pMD2. G (VSV-G envelope-expressing plasmid - 1 μg/flask), psPAX2 (lentiviral packaging plasmid - 1.5 μg/flask), and each CAR-vector plasmid (2.5 μg/flask) using Lipofectamine 3000 following the manufacturer’s instructions. The cell culture supernatant was collected every 24 hours and replaced with fresh medium during 72 hours, and the lentiviral particles were concentrated using Lenti-X concentrator reagent according to the manufacturer’s instructions. The lentiviral particles diluted in PBS were frozen at -80 °C, and the titer of the lentiviral particles was determined in Jurkat cells using a spinoculation protocol (850 × g for 65 min at RT) as previously reported (28). The formula {[(%GFP/100) × dilution × cell seeded]/final volume} was used to calculate the titer of lentiviral particles expressed as transducing units per mL (TU/mL).

### Generation of CAR-expressing Jurkat and NK-92 cells

Jurkat or NK-92 cells (1 × 10^5^ cells/250 µL/well) were seeded in 48-well plates containing appropriate medium, and cell transduction was performed using a spinoculation protocol (850 × g, at room temperature, for 65 minutes). Jurkat cells were modified with pLenti-mock, scFv3-, scFv12-, or scFvκ3-1-CAR at an MOI of 5 or with scFv5-CAR at an MOI of 10. NK-92 cells were modified with scFvκ3-1-CAR at an MOI of 10. Immediately after transduction, the modified cells were incubated in a humidified atmosphere with 5% CO_2_ at 37 °C for 72 hours. The transduction efficiency was evaluated using flow cytometry with GFP expression as a reporter. The cell concentration and viability were determined using propidium iodide (PI) (10 µg/mL) staining. Jurkat and NK-92 cells expressing CAR were expanded for 7 days and were enriched using fluorescence-activated cell sorting (FACS), which isolated a cell population expressing high levels of GFP. The sorted cells were expanded to create a cell stock containing Jurkat and NK-92 cells modified with CAR or pLenti-mock.

### Detection of CAR expression on the cell surface

The expression of scFv3-, scFv5-, and scFv12-CAR on the surface of Jurkat cells was detected by the presence of a Flag-tag in the N-terminus of the CAR. Cells modified with pLenti-Mock were used as a negative control. Jurkat cells (1 × 10^6^ cells/tube) were pelleted (300 × g, 10 min, 4 °C) and blocked with 0.5% BSA/PBS for 30 min. The cells were washed twice with PBS and incubated with an anti-FLAG monoclonal antibody (5 μg/mL; Sigma Aldrich) for 30 min. The cells were washed with PBS and incubated with a biotinylated goat secondary antibody. The detection of CAR was using streptavidin–phycoerythrin (PE) at a dilution of 1:100. After 30 min of incubation, the cells were washed and resuspended in PBS, and the percentage of cells positive for FLAG was measured by flow cytometry (Guava Easycyte Mini). All the incubations were performed on ice.

### Incubation of CAR-expressing Jurkat and NK-92 cells with *Candida* spp. or carbohydrates

Jurkat cells (2 × 10^5^/mL) or NK-92 cells (5 × 10^5^/mL) expressing CAR were dispensed into 96-well plates containing live or heat-killed yeast cells and hyphae that were cocultured with different ratios of target (T) to effector (E) cells. Jurkat cells were cocultured at a T:E ratio of 1:1 for heat-killed yeast cells and hyphae and 1:100, 1:200, and 1:400 for live yeast cells. The NK-92 cells were cocultured at a T:E ratio of 1:50, 1:100, or 1:200 relative to live yeast cells. In addition, Jurkat cells modified with or without CAR were incubated with mannan from *Saccharomyces cerevisiae* (1 μg/mL), beta-glucan (1 μg/mL), curdlan (1 μg/mL), a cell wall protein extract of *C. albicans* (1 μg/mL), phorbol 12-myristate 13-acetate (PMA) (50 ng/mL) plus ionomycin (1 μM), or phytohemagglutinin (PHA-L) (5 µg/mL) + PMA (50 ng/mL) as positive controls. Jurkat cells transduced with Mock were used as a negative control for cell activation. Cells were incubated in a humidified atmosphere with 5% CO_2_ at 37 °C for 24 hours and harvested to detect the expression of exhaustion and activation markers by flow cytometry, as described below. The levels of cytokines in the cell culture supernatants were measured by ELISA. The levels of IL-2 (BD OptiEIA Human IL-2 kit) and IFN-γ (BD OptiEIA Human IFN-γ kit) in the supernatant indicated the cell activation of Jurkat and NK-92 cells, respectively.

### Detection of exhaustion and activation markers on the cell surface and intracellular staining for phospho-ZAP-70 by flow cytometry

The expression of exhaustion markers (PD-1 and TIM-3) and activation markers (CD25 and CD69) was evaluated in Jurkat cells expressing or not expressing CAR. The heat-killed yeast or hyphae of *C. albicans* were cocultured with modified Jurkat cells, as described above, and the cells incubated with medium only were used as a negative control. The cocultures were incubated in a humidified atmosphere with 5% CO_2_ at 37

°C; after 24 h of incubation, the cells were collected to detect PD-1, TIM-3, CD25, or CD69. The cells were blocked with 1% BSA/PBS for 30 minutes and washed with PBS. The cells were incubated with anti-PD-1, anti-TIM-3, anti-CD25, or anti-CD69 monoclonal antibodies for 30 min, and washed. The percentage of positive cells was measured by flow cytometry (Guava Easycyte Mini). The incubation and washing steps were performed at 4 °C.

Jurkat cells expressing scFvκ3-1-CAR or Mock cells (2 × 10^5^ cells/mL) were seeded in a 96-well plate and incubated with medium alone, heat-killed yeast cells or with the hyphae of *C. albicans* at an E:T ratio of 1:1. After 10 min of incubation, the cells were harvested and pelleted (300 × g, 10 min, at 4 °C) before the addition of fixation/permeabilization solution, according to the manufacturer’s instructions. The cells were incubated with an anti-phospho-ZAP70 monoclonal antibody for 45 min and washed once before the percentage of cells positive for phosphor-ZAP-70 was determined by flow cytometry (Guava Easycyte Mini). The phosphorylation of ZAP-70 was induced with H_2_O_2_ (0.03%) as a positive control. All incubation and washing steps were performed at 4 °C.

### Fluorescence microscopy to monitor the redirection of modified cells to target *C. albicans*

Heat-killed yeast cells and hyphae (2 × 10^5^ cells/mL) were stained with Calcofluor white (100 µg/mL) for one hour and washed once before incubation with Mock or scFvκ3-1-CAR Jurkat cells (2 × 10^5^ cells/mL) in a 96-well plate. The coculture was incubated in a humidified atmosphere with 5% CO_2_ at 37 °C for 24 hours, and the targeting of the yeast cells and hyphae (blue) by modified Jurkat cells (green) was visualized under a fluorescence microscope (Leica) at 200× magnification. The interactions between NK-92 cells (5 × 10^5^ cells/ml) modified or not modified with scFvκ3-1-CAR and *C. albicans* were also analyzed by fluorescence microscopy. Live *C. albicans* yeast cells were incubated with modified NK-92 cells in a 96-well plate at ratios of target to effector of 1:50 and 1:100. After 6 hours of incubation, Calcofluor white (10 µg/mL) was added to the coculture for labeling of *C. albicans* for 15 minutes before visualization under a fluorescence microscope (Leica) at 200× magnification.

### Effects of the pharmacological inhibitors dasatinib and R406 on cell activation mediated by the scFvκ3-1-CAR

Jurkat cells expressing scFvκ3-1-CAR and Mock cells (2 × 10^5^ cells/mL) were seeded in a 96-well culture plate and treated with dasatinib (50 nM) and/or R406 (2 µg/mL) for 3 hours. Then, the cells were cocultured with heat-killed yeasts and hyphae of *C. albicans* at an E:T ratio of 1:1 in a humidified atmosphere with 5% CO_2_ at 37 °C, and after 24 hours of incubation, the plate was centrifuged (300 × g, 10 min, 25 °C), and the levels of IL-2 in the cell culture supernatants were measured by ELISA

### Apoptosis analysis using Annexin V staining

Jurkat cells modified with pLenti-Mock or scFvκ3-1-CAR (2 × 10^5^/mL) were cocultured with heat-killed yeast or hyphae of *C. albicans* in a 96-well cell culture plate at an E:T ratio of 1:1. Arsenic trioxide (As_2_O_3_, 12 µM) was used as a positive control to induce apoptosis, and the cells incubated only with medium were considered a negative control. After 24 hours of incubation in a humidified atmosphere with 5% CO_2_ at 37 °C, 10x binding buffer was added to the cell culture prior to incubation with PE-conjugated Annexin V (2 µL/well/200 µL). After 30 minutes of incubation at 37 °C, the cells were harvested, and the percentage of Mock and scFvκ3-1-CAR Jurkat cells positive for Annexin V was determined by flow cytometry.

### Detection of CD107a expression by flow cytometry

Nontransduced NK-92 cells and NK-92 cells expressing scFvκ3-1-CAR (5 × 10^5^/mL) were seeded in a 96-well microplate and incubated with live *C. albicans* yeast cells at a T:E ratio of 1:100. Thereafter, monensin (2 µM) and a FITC-conjugated anti-human CD107a monoclonal antibody were added to the cells at the beginning of the coculture. PMA (50 ng/mL) plus ionomycin (1 μM) was used as a positive control to induce the release of cytotoxic granules, and NK-92 cells incubated with medium alone were used as a negative control. After 6 hours of coculture in a humidified atmosphere with 5% CO_2_ at 37 °C, the cells were collected, and the expression of CD107a was detected by flow cytometry.

### *In vitro* anti-*Candida* spp. activity of NK-92 cells expressing scFvκ3-1-CAR

For the CFU assay, NK-92 cells (5 × 10^5^ cells/mL) modified with or without scFvκ3-1-CAR were cocultured with live yeast cells of *C. albicans, C. tropicalis* or *C. glabrata* at a T:E ratio of 1:100. The cocultures were incubated for 24 hours in a humidified atmosphere with 5% CO_2_ at 37 °C, and the samples were homogenized, diluted in sterile PBS and plated on Sabouraud Dextrose agar medium supplemented with kanamycin (50 µg/mL). The plates were incubated at 30 °C for 24–36 h, and the number of colonies was determined to express the fungal burden as CFU/mL. The growth of yeast cells in the absence of NK-92 cells was taken as a negative control.

To perform the XTT assay, NK-92 cells expressing scFvκ3-1-CAR (5 × 10^5^ cells/mL) were incubated with live *C. albicans* yeast cells in a 96-well plate at T:E ratios of 1:25 and 1:50. After 4 and 6 hours of incubation in a humidified atmosphere with 5% CO_2_ at 37 °C, the plates were centrifuged (800 × g, 10 min, RT), and the NK-92 cells were lysed with sterile distilled water for 45 minutes. The plates were centrifuged (800 × g, 10 min, RT), and the pelleted *C. albicans* was incubated with an XTT solution (0.25 mg/mL) supplemented with menadione (1 µL). After 2 hours of incubation at 37 °C, the absorbance at 450 nm was recorded by a spectrophotometer. The absorbance at 690 nm was also obtained as a reference. The calculation for hyphal damage was as follows: hyphal damage [%] = (1 - X/C) × 100, where X represents the absorbance of the experimental wells and C represents the absorbance of the control wells with hyphae only.

### NSG mice infected with *C. albicans* and treated with scFvκ3-1-CAR-NK-92 cells

The kinetics of the dissemination of *C. albicans* were evaluated in female (*n* = 10) and male (*n* = 10) NSG mice infected with 1 × 10^4^ *C. albicans* yeast cells (50 µL) intravenously via the retro-orbital route. After 6, 24, and 72 hours of infection, the animals were euthanized to obtain the liver, kidney, lung, and spleen, which were homogenized in 1 mL of cold sterile PBS buffer (pH 7.2). The tissue suspensions were plated on Sabouraud dextrose agar media supplemented with kanamycin (50 µg/mL). After 24–36 h of incubation at 30 °C, the number of colonies was determined, and the fungal burden was expressed in CFU/g.

Male NSG mice were divided into three groups: Untreated (*n* = 10), NK-92 cells (*n* = 17), and scFvκ3-1-CAR NK-92 cells (*n* = 22). All mice were infected intravenously via the retro-orbital route with 1 × 10^4^ yeasts of *C. albicans* diluted in 50 µL of PBS. After 3 hours of infection, a suspension of NK-92 cells expressing or not expressing scFvκ3-1-CAR (5 × 10^6^ cells/animal) diluted in 100 µL of PBS was infused via the lateral tail vein. Untreated mice received only vehicle-treated PBS via the lateral tail vein. Mice were euthanized at 24 hours postinfection, the lung, liver, and kidney were collected aseptically, and 1 mL of cold sterile PBS buffer (pH 7.2) was added to each. The organs were immediately weighed and homogenized. Then, the tissue suspension was diluted in PBS and plated on Sabouraud dextrose agar medium supplemented with 50 µg/mL kanamycin. The plates were incubated at 30 °C for 24–36 h, and the fungal burden was expressed in CFU/g. Three biological replicates were performed.

### scFv κ3-1 protein expression and purification

The *E. coli* strain NiCo21(DE3) was transformed by heat shock with a plasmid containing the scFv κ3-1 sequence with a His-tag at its N-terminus (Fig. S2A). After colony growth on LB agar medium containing carbenicillin (50 µg/mL), scFv κ3-1 protein expression was induced using the autoinduction method with terrific broth (TB) medium. Bacteria were cultivated at 37 °C and agitated at 200 rpm until the optical density (O.D.) reached 0.05-0.1. The induction was then carried out overnight at 22 °C at 200 rpm. After induction, the cells were harvested by centrifugation (4000 RPM, 20 min, 4 °C), and the cell pellets were resuspended in HBS buffer. Next, the samples were homogenized with 2 protease inhibitor cocktail tablets (Roche). The cells were lysed by sonication and centrifuged at 18000 rpm at 4 °C for 30 minutes to separate the insoluble and soluble fractions. The supernatants containing soluble protein were used for the purification steps.

The scFv κ3-1 protein was purified by nickel affinity (Ni-NTA) chromatography, with elution at imidazole concentrations of 30 mM, 50 mM, 100 mM, 150 mM, and 300 mM in HBS buffer. The purification efficacy was assessed by SDS‒PAGE, analyzing samples from each imidazole concentration step during elution (Fig. S2A). The gel was then stained with Coomassie blue. The expected molecular weight of the scFv κ3-1 protein is 28 kDa. The corresponding band was eluted with 100 mM imidazole (Fig. S2B), and this fraction was concentrated using a 10 kDa spin filter concentrator (Vivaspin 2-10000 MWCO) and subjected to size exclusion chromatography (SEC) using HPLC (AKTA). The column used was a Superdex 75 16/600 column. The SEC process yielded 47 fractions of 2 mL each. Fractions 11 to 20 were analyzed by SDS‒PAGE, and fractions 17, 18, 19, and 20 were considered the corresponding fractions of the scFv κ3-1 protein, exhibiting a single relatively intense band at approximately 28 kDa (Fig. S2C, highlighted in red). The fractions were pooled and concentrated to 500 µL each with a 10 kDa MWCO spin concentrator column. Sodium azide 0.02% (w/v) was added to the samples as a preservative. The samples were snap-frozen in liquid nitrogen and stored at –20 °C.

### NMR analysis of the scFv κ3-1 protein

Nuclear magnetic resonance (NMR) analysis of the scFv κ3-1 protein was performed. The sample was spiked with 10% D_2_O to reference the spectrum to a deuterium lock. The NMR analysis was conducted on a 600 MHz Bruker Avance III spectrometer, collecting a 1D proton spectrum with pre-saturation for water suppression (zgesgp, 2048 scans) at 298 K.

### Glycan microarray analysis

The binding specificity of the His-tagged scFv κ3-1 protein was evaluated using two types of carbohydrate microarrays: (1) a microarray designated ‘Fungal and Bacterial Polysaccharide Array’ featuring 19 saccharides (polysaccharides or glycoproteins) and one neoglycolipid (NGL) derived from chitin hexasaccharide (Table S2); and (2) an NGL-based sequence-defined glycan screening array composed of 672 lipid-linked oligosaccharide probes, of mammalian and non-mammalian type, essentially as previously described (49). The complete list of probes and their sequences can be found in Table S3.

The microarray analyses were performed essentially as described previously (80). In brief, microarray slides were blocked for 1 hour using a blocker/diluent solution comprising 10 mM HEPES (pH 7.4), 150 mM NaCl, 1% (w/v) bovine serum albumin (Sigma 7030), 0.02% Casein blocker (Pierce) and 5 mM CaCl_2_. The scFv κ3-1 protein was added at a concentration of 150 µg/mL and incubated for 90 minutes, followed by incubation with mouse monoclonal anti-His and biotinylated anti-mouse IgG antibodies (both from Sigma and overlaid at 10 µg/mL). Biotinylated plant lectin ConA (Vector Laboratories) was used at 2 µg/mL. The final detection was performed with a 30-minute overlay of streptavidin-Alexa Fluor 647 (Molecular Probes) at 1 µg/mL. All steps of the analyses were performed at room temperature. Details of the glycan library, the generation of the microarrays, imaging, and data analysis are in the Supplementary glycan microarray document (Tables S2 and S3) in accordance with the Minimum Information Required for A Glycomics Experiment (MIRAGE) guideline for reporting glycan microarray-based data (81).

### Statistical analysis

Statistical analysis was conducted using Prism 9.0 (GraphPad Software). The Shapiro–Wilk test was used to assess the normality of the data, and the homogeneity of variances was evaluated using Bartlett’s test for experiments with three or more groups. For normal distribution experiments, variance analysis (ANOVA) was applied with Tukey’s multiple comparisons test to assess the significance of differences between groups. The Kruskal‒Wallis test was used with Dunn’s multiple comparisons test for experiments with a nonnormal distribution. Statistical significance was set at p < 0.05. The results are reported as either the mean ± standard deviation (SD) or median with interquartile range.

## Additional Information and Declarations

### Author Contributions

G.Y.C. and T.A.S. conceived and designed the experiments, performed the experiments, analyzed the data, prepared figures and/or tables, authored or reviewed drafts of the article, and approved the final draft.

J.G.G., M.P.M., P.K.M.O.B., B.S., A.D.M, Y.L., S.J.M., and T.F. performed the experiments, analyzed the data, prepared figures and/or tables, and approved the final draft.

D.S., P.V.B.P., T.F.R., and G.H.G. performed the experiments and approved the final draft.

### Animal Ethics

All animal experiments were approved by the Committee on Ethics in Animal Research of Ribeirão Preto Medical School (protocol 071/2021). *C. auris* (clinical isolates 467, 468, and 469) were available to make a library of G.H.G.’s lab, and the Committee of Ethics of the University of São Paulo, Campus of Ribeirão Preto, Brazil (Permit Number: 08.1.1277.53.6; Studies on the interaction of fungal pathogens with animals) approved all protocols to work with clinical isolates.

