## Supplementary Information for "Enhanced antifungal activity of NK-92 cells against *Candida albicans* mediated by a mannan-specific chimeric antigen receptor"

### Supplementary Figure S1

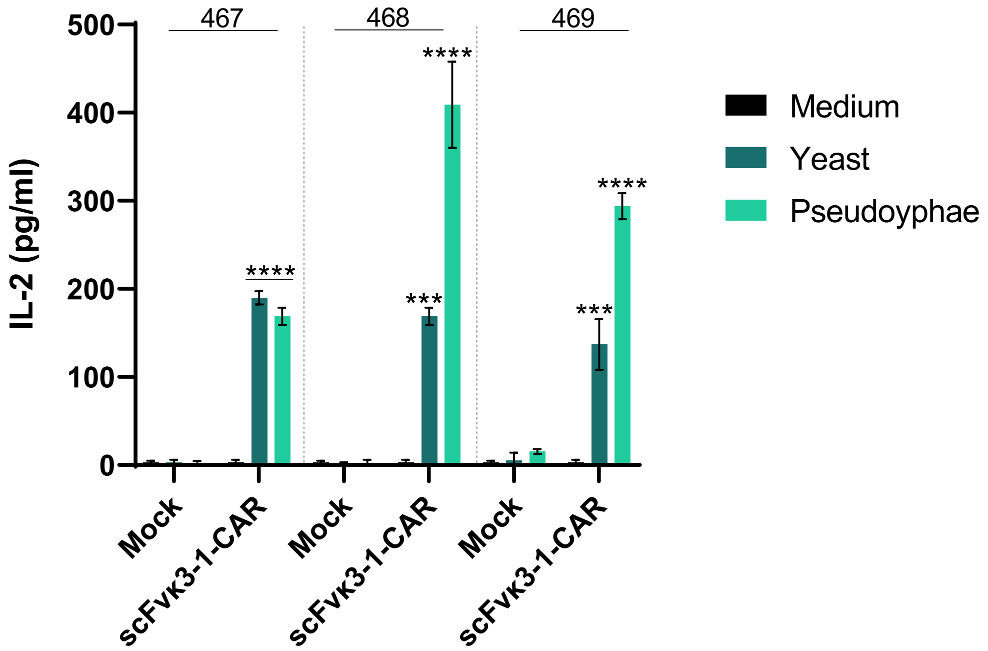


**Figure S1. scFvκ3-1-CAR mediates Jurkat cell activation in the presence of *Candida auris* clinical isolates.** The activation assay was performed with Jurkat cells expressing Mock or scFvκ3-1-CAR ($2 \times 10^5$ cells/mL). The cells were cocultured with heat-killed yeast and pseudohyphae of *Candida auris* clinical isolates 467, 468, and 469 (1:1 ratio) in 96-well plates. After 24 hours, the cell supernatant was collected and used to measure IL-2 levels by ELISA (pg/mL) ($n = 3$). The groups were compared to the "medium" group. The values are expressed as the mean ± SD. The experiment was performed in triplicate. *** $p < 0.001$, **** $p < 0.0001$.

**Supplementary Table S1. Screening for cell activation triggered by Mock, scFv5-CAR and scFvκ3-1-CAR constructs in the presence of *Candida* species and other fungal genera.**

| | Mock | | scFv5-CAR | | scFvκ3-1-CAR | |
|---|---|---|---|---|---|---|
| | IL-2 (pg/mL) | *p* value | IL-2 (pg/mL) | *p* value | IL-2 (pg/mL) | *p* value |
| *Candida guilliermondii* (yeast) | 0.8612 ± 1.218 | 0.5000 | 0.00 | NA | 2.105 ± 3.646 | 0.5421 |
| *Candida guillermondii* (pseudohyphae) | 0.00 | NA | 0.00 | NA | 3.828 ± 3.265 | 0.8938 |
| *Candida krusei* (yeast) | 0.00 | NA | 0.8193 ± 1.802 | 0.8005 | 0.00 | NA |
| *Candida krusei* (pseudohyphae) | 0.00 | NA | 1.419 ± 2.707 | 0.4052 | 0.00 | NA |
| *Candida parapsilosis* (yeast) | 0.8612 ± 1.218 | 0.9291 | 0.00 | NA | 2.297 ± 2.297 | 0.4132 |
| *Candida parapsilosis* (pseudohyphae) | 3.158 ± 4.466 | 0.4799 | 0.00 | NA | 1.148 ± 1.519 | 0.2125 |
| *Aspergillus fumigatus* | 3.839 ± 2.921 | 0.6300 | 6.888 ± 9.679 | 0.8149 | 1.468 ± 0.8525 | 0.4906 |
| *Cryptococcus gattii* | 0.00 | NA | 5.380 ± 1.541 | 0.5347 | 0.00 | NA |
| *Cryptococcus neoformans* | 0.6745 ± 1.168 | 0.4226 | 17.15 ± 3.530 | 0.1379 | 0.00 | NA |
| *Cryptococcus neoformans* (acapsular mutant) | 0.00 | NA | 6.220 ± 5.578 | 0.6916 | 0.00 | NA |
| *Rhizopus oryzae* | 0.00 | NA | 1.849 ± 1.771 | 0.1279 | 0.00 | NA |

*NA* not applicable. *p* value is for comparisons between each modified cell group co-cultured with the target vs without the target.

### Supplementary Figure S2

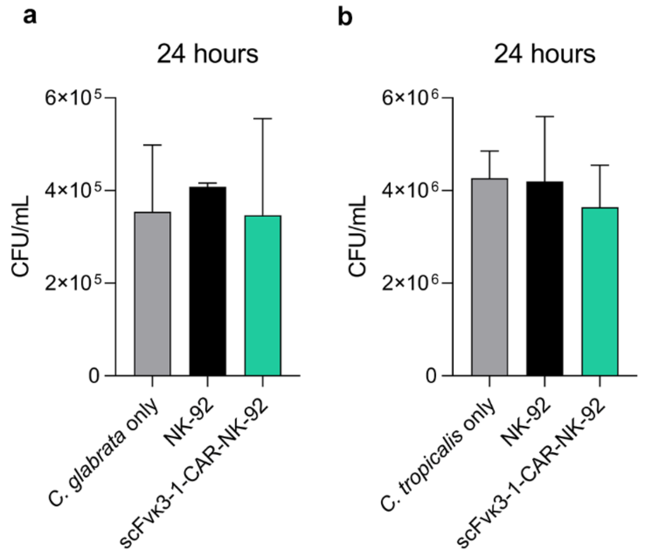


**Figure S2. scFvκ3-1-CAR-NK-92 cells did not affect *C. glabrata* and *C. tropicalis* growth.** NK-92 cells modified or not with scFvκ3-1-CAR (5 × $10^5$ cells/mL) were cocultivated with (**a**) *C. glabrata* (*n* = 3–4) and (**b**) *C. tropicalis* (*n* = 4) in a 96-well plate at a ratio of 1:100. After 24 hours of incubation, the cell suspensions were diluted in PBS and plated to perform the CFU assay, and the results were presented as CFU/mL. *C. glabrata* and *C. tropicalis* alone were used as controls.

### Supplementary Figure S3

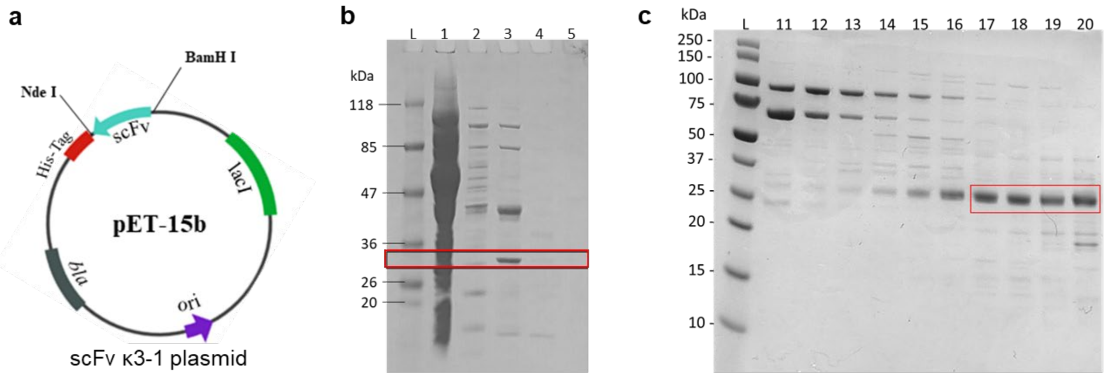


**Figure S3. Expression and purification of the scFv κ3-1 protein.** (**a**) Scheme of the plasmid designed for transformation of the *E. coli* strain NiCo21(DE3), which contains the scFv κ3-1 coding sequence and an N-terminal His-tag. (**b**) SDS–PAGE analysis of scFv κ3–1 protein expression after elution with different concentrations of imidazole. L. Molecular weight protein marker (kDa); 1. Imidazole (30 mM); 2. 50 mM; 3. 100 mM; 4. 150 mM; 5. 300 mM. The red rectangle indicates the expected size of the scFv κ3-1 protein (28 kDa). (**c**) SDS–PAGE analysis of the scFv κ3-1 protein after size exclusion chromatography, which revealed fractions 11 to 20. L. Molecular weight protein marker (kDa). The red rectangle indicates the expected size of the scFv κ3-1 protein (17, 18, 19, 20).

### Supplementary Figure S4

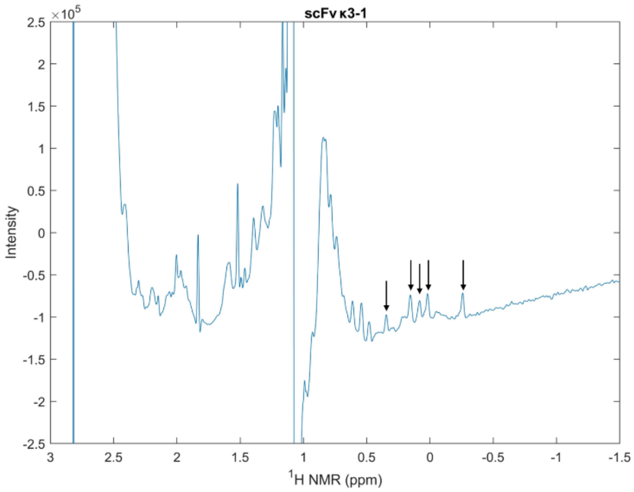


**Figure S4. 1D NMR spectrum of the scFv κ3-1 protein**. NMR analysis was conducted on a 600 MHz Bruker Avance III spectrometer. The black arrows indicate methyl peaks around the 0-ppm region.

**Supplementary Table S2. Saccharide probes included in the Fungal and Bacterial Polysaccharide Array.**

| Position | Probe[a] | Predominant oligosaccharide sequences where known[b] |
|---|---|---|
| 1 | Dextran *L. mesenteroides* | α1,6-Glc |
| 2 | Pullulan *P. pullulans* | Mixed α1,4/α1,6-Glc |
| 3 | Curdlan[c] *Agrobacterium* sp. | β1,3-Glc |
| 4 | NSG *S. cerevisiae* | Linear β1,3-Glc backbone with occasional monoglucosyl β1,6-Glc branches |
| 5 | PGG *S. cerevisiae* | |
| 6 | Lentinan *L. edodes* | |
| 7 | Grifolan *G. frondosa* | β1,3-Glc backbone with highly ramified oligomeric branches |
| 8 | Barley β-glucan | Mixed β1,3/β1,4-Glc |
| 9 | Oat β-glucan | Mixed β1,3/β1,4-Glc |
| 10 | Lichenan | Mixed β1,3/β1,4-Glc |
| 11 | Pustulan *U. papullosa* | β1,6-Glc |
| 12 | Mannan *S. cerevisiae* | α1,6-Man backbone with oligomeric α1,2-, α1,3-Man branches [42] |
| 13 | *N*-Mannoprotein *C.albicans* | α1,6-Man backbone with oligomeric α1,2-, α1,3-, and β-1,2-Man branches [42-44] |
| 14 | Mannoprotein *A. fumigatus* | Mannose-rich with β1,5-galactofuranose moieties [82,83] |
| 15 | Lipomannan *M. tuberculosis* | Linear α1,6-Man backbone with monomannosyl α1,2-Man branches [84] |
| 16 | Lipoarabinomannan *M. tuberculosis* | Linear α1,6-Man backbone with monomannosyl α1,2-Man branches and α1,5-Ara polymer branched at certain positions with α1-3,1-5-Ara residues, which in turn are terminated by β1,2-Ara and capped by α1,2-Man units [84] |
| 17 | Lipoarabinomannan *M. smegmatis* | Linear α1,6-Man backbone with monomannosyl α1,2-Man branches and α1,5-Ara polymer branched at certain positions with α1-3,1-5-Ara residues, which in turn are terminated by β1,2-Ara and capped by phospho inositol [84] |
| 18 | Native *O*-glycoprotein *M. tuberculosis* | α1,2-Man [84] |
| 19 | Glucurono-XyloMannan[d] *T. fuciformis* | α1,3-Man with Xyl, GlcA and Fuc branches |
| 20 | GN6-AO[e] | GlcNAcβ-4GlcNAcβ-4GlcNAcβ-4GlcNAcβ-4GlcNAcβ-4GlcNAc-AO [80] |

[a] Unless otherwise indicated the saccharide probes are polysaccharides: NSG, Neutral soluble β-glucan; PGG, Poly-(1,6)-D-glucopyranosyl-(1,3)-D-glucopyranose

[b] Glc, Glucose; Man, Mannose; Gal, Galactose; Ara, Arabinose; Xyl, xylose; GlcA, Glucuronic acid; Fuc, fucose; GlcNAc, *N*-acetylglucosamine.

[c,d] Curdlan polysaccharide was solubilized in 50mM NaOH and Glucurono-XyloMannan in 150 mM NaCl, prior printing.

[e] GN$_6$-AO, neoglycolipid (NGL) probe prepared from reducing hexasaccharide of chitin (β1,4-linked *N*-acetylglucosamine, GlcNAc) by oxime ligation with an aminooxy (AO) functionalised DHPE [80].

**Supplementary Table S3. Comprehensive Screening Array**

Please find this table attached separately (Excel file).

**Supplementary Figure S5**

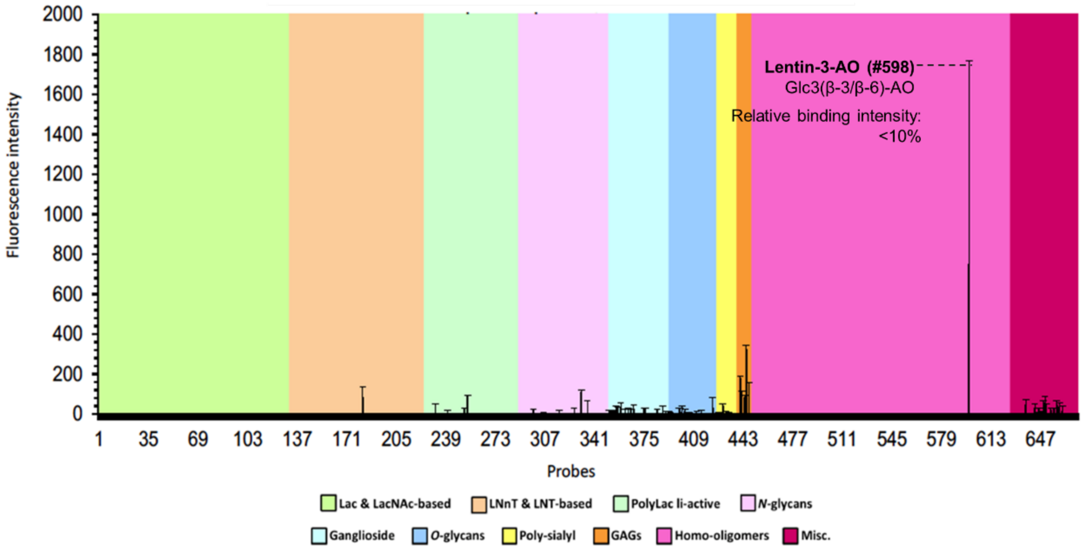


**Figure S5. Comprehensive sequence-defined glycan screening analysis of scFv κ3-1 binding.** The scFv κ3-1 protein (150 µg/mL) was used for comprehensive sequence-defined glycan screening analysis with 672 glycan probes. The binding of the scFv κ3-1 protein was determined by fluorescence intensity. The glycan probes are grouped according to their backbone-type sequences as annotated by the following colored panels: disaccharide-based probes, lactose (Lac) and Nacetyllactosamine (LacNAc); tetrasaccharide-based probes, lacto-N-neo-tetraose (LNnT) and lacto-N-tetraose (LNT); poly-Nacetyllactosamine (PolyLacNAc); N-glycans; gangliosides; O-glycan-related probes; polysialyl probes; glycosaminoglycans (GAGs); homooligomers of glucose and other monosaccharides; and other nonclassified sequences (miscellaneous, Misc). The complete list of probes and their sequences are provided in Table S3.
